# Hepatitis C Virus Drugs Simeprevir and Grazoprevir Synergize with Remdesivir to Suppress SARS-CoV-2 Replication in Cell Culture

**DOI:** 10.1101/2020.12.13.422511

**Authors:** Khushboo Bafna, Kris White, Balasubramanian Harish, Romel Rosales, Theresa A. Ramelot, Thomas B. Acton, Elena Moreno, Thomas Kehrer, Lisa Miorin, Catherine A. Royer, Adolfo García-Sastre, Robert M. Krug, Gaetano T. Montelione

**Author notes:** These two investigators made equal contributions to this project (Khushboo Bafna, Kris White).

## Abstract

Effective control of COVID-19 requires antivirals directed against SARS-CoV-2 virus. Here we assess ten available HCV protease inhibitor drugs as potential SARS-CoV-2 antivirals. There is a striking structural similarity of the substrate binding clefts of SARS-CoV-2 M^pro^ and HCV NS3/4A proteases, and virtual docking experiments show that all ten HCV drugs can potentially bind into the M^pro^ binding cleft. Seven of these HCV drugs inhibit SARS-CoV-2 M^pro^ protease activity, while four dock well into the PL^pro^ substrate binding cleft and inhibit PL^pro^ protease activity. These same seven HCV drugs inhibit SARS-CoV-2 virus replication in Vero and/or human cells, demonstrating that HCV drugs that inhibit M^pro^, or both M^pro^ and PL^pro^, suppress virus replication. Two HCV drugs, simeprevir and grazoprevir synergize with the viral polymerase inhibitor remdesivir to inhibit virus replication, thereby increasing remdesivir inhibitory activity as much as 10-fold.

**Highlights:**

- Several HCV protease inhibitors are predicted to inhibit SARS-CoV-2 M^pro^ and PL^pro^.
- Seven HCV drugs inhibit M^pro^ enzyme activity, four HCV drugs inhibit PL^pro^.
- Seven HCV drugs inhibit SARS-CoV-2 replication in Vero and/or human cells.
- HCV drugs simeprevir and grazoprevir synergize with remdesivir to inhibit SARS-CoV-2.

**eTOC blurb:** Bafna, White and colleagues report that several available hepatitis C virus drugs inhibit the SARS-CoV-2 M^pro^ and/or PL^pro^ proteases and SARS-CoV-2 replication in cell culture. Two drugs, simeprevir and grazoprevir, synergize with the viral polymerase inhibitor remdesivir to inhibit virus replication, increasing remdesivir antiviral activity as much as 10-fold.

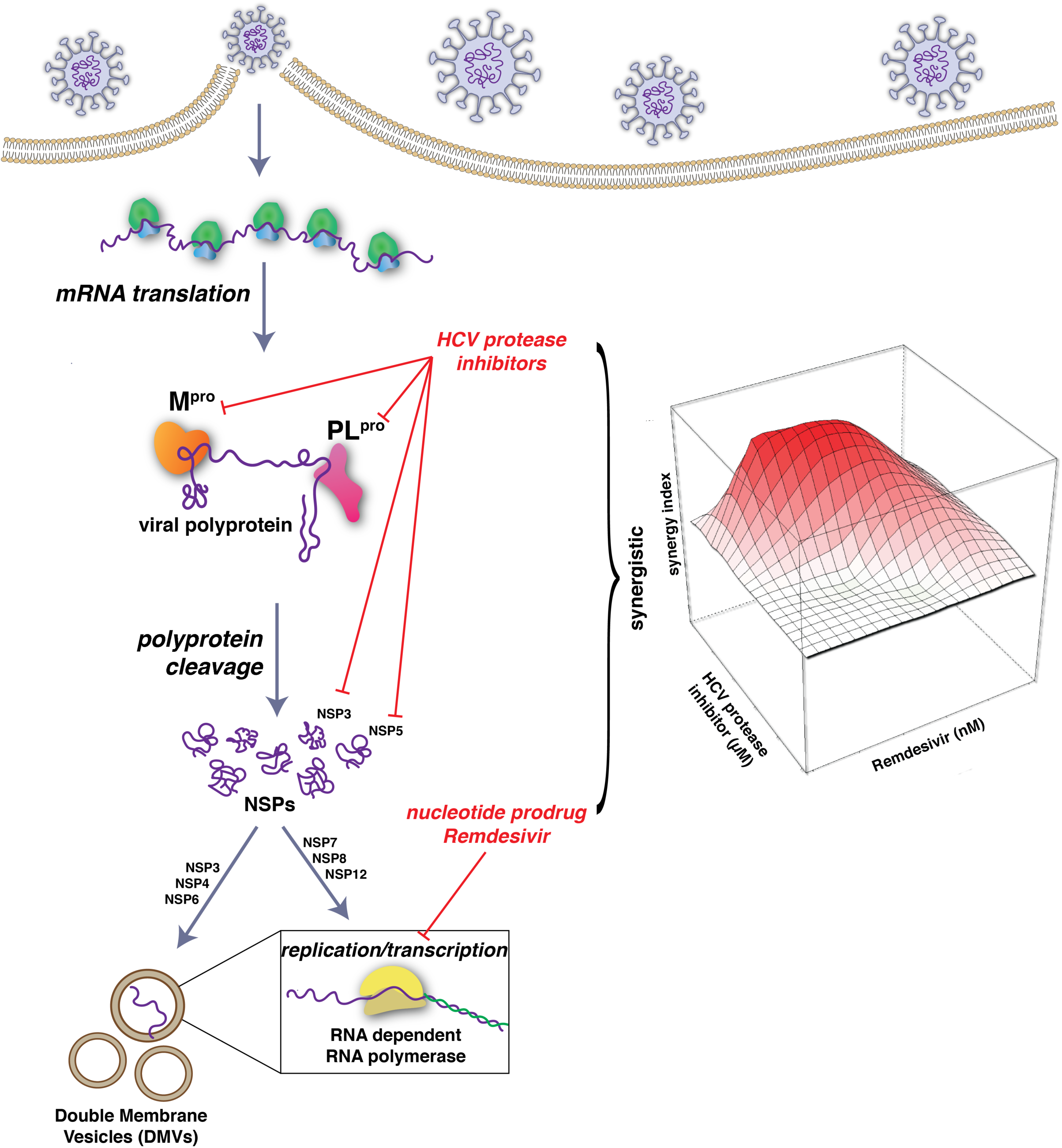

## Introduction

The COVID-19 pandemic has caused more than one million deaths worldwide and crippled the global economy. Effective control of the SARS-CoV-2 coronavirus that causes COVID-19 requires antivirals, especially until safe and effective vaccines are available. Considering the urgency to identify effective antiviral drugs, and the usually lengthy process involved in approving candidate drugs for human use, our goal is to identify existing drugs already approved for use in humans that can be repurposed as safe and effective therapeutics for treating COVID-19 infections, and which may also be useful as lead molecules for novel drug development.

SARS-CoV-2 is an enveloped RNA virus which causes COVID-19 (Wu et al., 2020). Its genome is comprised of a single, large positive-sense single-stranded RNA, which is directly translated by host cell ribosomes. The SARS-CoV-2 genome encodes 4 structural proteins, 16 non-structural proteins (NSPs) which carry out crucial intracellular functions, and 9 accessory proteins (Gordon et al., 2020; Wu et al., 2020). Many of these proteins, and their host binding partners, are potential targets for development of antivirals for SARS-CoV-2. For example, the repurposed drug remdesivir, which inhibits the viral RNA-dependent RNA polymerase, is the current FDA-approved antiviral standard of care for COVID-19 (Eastman et al., 2020; Pan et al., 2020).

Translation of the viral genomic RNA results in the biosynthesis of two polyproteins that are processed into the 16 separate NSPs by two virus-encoded cysteine proteases, the papain-like protease (PL^pro^) and a 3C-like protease (3CL^pro^). The latter is also referred to as the main protease (M^pro^). M^pro^ and PL^pro^ are essential for the virus life cycle, specifically including the production of a functional viral RNA polymerase, and are hence attractive targets for antiviral development. M^pro^ cleavages generate several NSPs including the three subunits nsp7, nsp8 and nsp12 of the viral RNA polymerase complex, as well as nsp4 and nsp6 integral membrane proteins (Peng et al., 2020). PL^pro^ cleavages generate integral membrane nsp3 protein. The nsp3 – nsp4 – nsp6 complex is required for forming replication organelles, also known as double membrane vesicles, that are required for the function of the viral RNA polymerase (Angelini et al., 2013; Oudshoorn et al., 2017; Snijder et al., 2020; Wolff et al., 2020a; Wolff et al., 2020b). Considering that both M^pro^ and PL^pro^ generate either the RNA polymerase itself, or the replication organelles required for polymerase function, we reasoned that inhibitors of one or both of these proteases might be synergistic with inhibitors of the viral polymerase, such as remdesivir.

We observed that the substrate binding cleft and active site of the SARS-CoV-2 M^pro^ has remarkable structural similarity with the active site of the hepatitis C virus (HCV) NS3/4A protease, suggesting that drugs that inhibit the HCV protease might also inhibit SARS-CoV-2 M^pro^ (Bafna et al., 2020). Consistent with this hypothesis, recent studies have reported that one of these available HCV drugs, boceprevir, inhibits M^pro^ proteolytic activity and SARS-CoV-2 replication in Vero cells (Anson et al., 2020; Fu et al., 2020; Ma et al., 2020). Several other HCV protease inhibitors have also been reported to inhibit M^pro^ proteolytic activity to various extents (Anson et al., 2020; Fu et al., 2020; Ma et al., 2020), while other studies report that some of these same HCV protease inhibitors did not significantly inhibit M^pro^ (Fu et al., 2020).

In this study we assess the abilities of ten available HCV protease inhibitors to suppress the replication of SARS-CoV-2. Virtual docking experiments predict that all ten HCV drugs can bind snuggly into the M^pro^ binding cleft with docking scores comparable to a known M^pro^ inhibitor, suggesting that any of these ten HCV drugs are potential inhibitors of M^pro^. Surprisingly, we found that some of these HCV drugs also dock well into the PL^pro^ substrate binding cleft. Seven of these HCV drugs inhibit both SARS- CoV-2 M^pro^ protease activity, and SARS-CoV-2 virus replication in Vero and/or human 293T cells expressing the SARS-CoV-2 ACE2 receptor. Three of these seven also inhibit PL^pro^ protease activity. Consequently, HCV drugs that inhibit M^pro^, or inhibit both M^pro^ and PL^pro^, can suppress SARS-CoV-2 virus replication, v*iz,* boceprevir (BOC), narlaprevir (NAR), vaniprevir (VAN), telaprevir (TEL), simeprevir (SIM), grazoprevir (GRZ), and asunaprevir (ASU).

Further, we demonstrate that two HCV drugs which inhibit the proteolytic activities of both M^pro^ and PL^pro^, simeprevir and grazoprevir, act synergistically with remdesivir to inhibit SARS-CoV-2 virus replication, thereby increasing remdesivir antiviral activity as much as 10-fold. In contrast, boceprevir, which inhibits M^pro^ but not PL^pro^, acts additively, rather than synergistically with remdesivir to inhibit virus replication. Simeprevir provides synergy with remdesivir at significantly lower concentrations than grazoprevir. Our results suggest that the combination of a HCV protease inhibitor with a RNA polymerase inhibitor could potentially function as an antiviral against SARS-CoV-2. In particular, the combination of two FDA-approved drugs, namely, simeprevir and remdesivir, could potentially function as a therapeutic for COVID-19 while more specific SARS-CoV-2 antivirals are being developed. More generally, our results strongly motivate further studies of the potential use of M^pro^ and/or PL^pro^ protease inhibitors in combination with RNA polymerase inhibitors as antivirals against SARS-CoV-2.

## Results

### Similarity of the substrate-binding clefts and active sites of SARS-CoV-2 M^pro^ and HCV protease NS3/4A

The SARS-CoV-2 main protease (M^pro^) is a 67.6 kDa homodimeric cysteine protease with three domains (Jin et al., 2020; Zhang et al., 2020b). Domains I and II adopt a double β-barrel fold, with the substrate binding site located in a shallow cleft between two antiparallel β-barrels of Domain I and II (Figure 1A). C-terminal domain III stabilizes the homodimer. The fold architecture of domains I and II are similar to picornavirus cysteine proteases and chymotrypsin serine proteases (Anand et al., 2002; Gorbalenya et al., 1989). A three-dimensional structural similarity search of the Protein Data Bank using the *DALI* program (Holm and Sander, 1993, 1999), with domains I and II (excluding domain III) of the SARS-CoV-2 M^pro^ as the three-dimensional structural query, identified several proteases, including the HCV NS3/4A serine protease, as structurally similar. These two enzymes have a structural similarity Z score (Holm and Sander, 1993, 1999) of +8.4, and overall backbone root-mean-squared deviation for structurally similar regions of ∼ 3.1 Å. The HCV serine protease also has a double β- barrel fold, with relative orientations of several secondary structure elements similar to those of the SARS-CoV-2 M^pro^ cysteine protease, and a substrate binding site located in a shallow cleft between its two six- to eight-stranded antiparallel β-barrels (Figure 1B). Superimposition of the analogous backbone structures of these two proteases results in superimposition of their substrate binding clefts and their active-site catalytic residues, His41 / Cys145 of the SARS-CoV-2 M^pro^ cysteine protease and His57 / Ser139 of the HCV NS3/4A serine protease (Figure 1C). Because of these structural similarities, we proposed that some HCV protease inhibitors might bind well into the substrate binding cleft of the SARS-CoV-2 M^pro^ and inhibit virus replication.

**Fig. 1.**
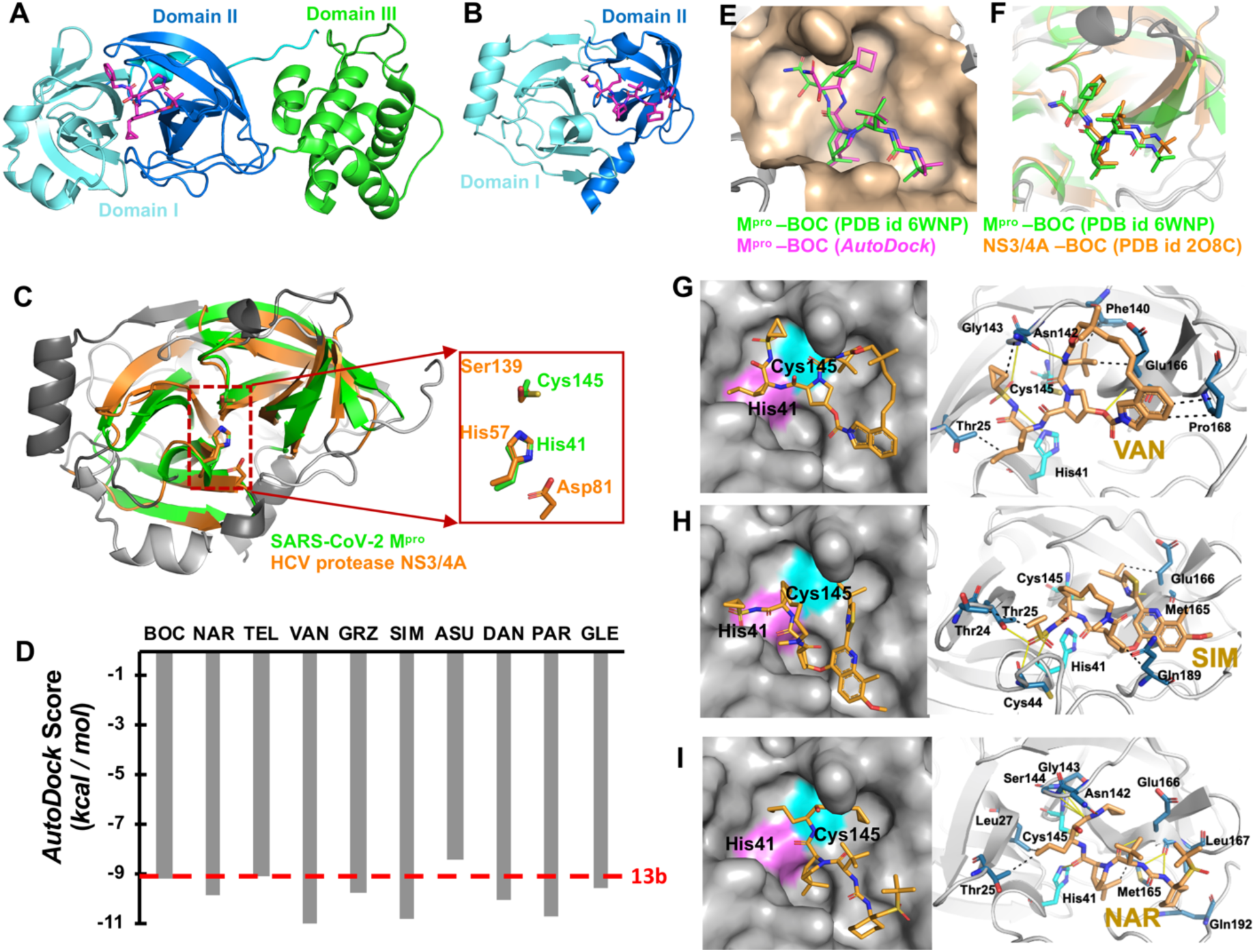
The substrate binding cleft and active site of SARS-CoV-2 M^pro^ and HCV protease NS3/4A have remarkable structural similarity. (A) SARS-CoV-2 M^pro^ (PDB 6Y2G) and (B) HCV protease NS3/4A (PDB 2P59) both have a double β-barrel fold architecture, with a substrate binding site located in a shallow cleft between two the antiparallel β-barrels (shown in cyan and blue). Only one protomer is shown for both structures. The a-helical dimerization domain III of M^pro^ is shown in green. The structures of bound inhibitors in these crystal structures are illustrated as magenta sticks. (C) The backbone structure of the SARS-CoV-2 M^pro^ (green) is superimposed on the backbone structure of HCV protease NS3/4A (orange). The regions identified by DALI as structurally-analogous are shown in color (green and orange), and the regions that are not structurally-analogous are shown in gray. This superimposition of backbone atoms results in superimposition of the catalytic residues Cys145 and His41 of the SARS-CoV-2 M^pro^ with Ser139 and His57 of HCV protease. Asp81 of the HCV protease catalytic triad is also shown. (D) *AutoDock* docking scores for 10 HCV protease inhibitors in the substrate binding cleft of SARS-CoV-2 M^pro^. The docking score for M^pro^ inhibitor 13b (*AutoDock* score = -9.03 kcal/mol) is also shown as a horizontal red dashed line. (E) Comparison of the boceprevir (BOC) binding pose in best-scoring *AutoDock* complex (magenta) with the X-ray crystal structure of BOC-M^pro^ complex (Anson and Mesecar, 2020) (green, PDB id 6WNP). (F) Comparison of BOC binding poses in X-ray crystal structures complexes with HCV NS3/4A protease (orange, PDB id 2OC8) and SARS-CoV-2 M^pro^ protease (green, PDB id 6WNP). (G - I) Best-scoring *AutoDock* poses for complexes of vaniprevir (VAN), simeprevir (SIM), and narlaprevir (NAR) with M^pro^. Residues forming hydrogen bonds (yellow solid line) and hydrophobic interactions (black dash) are labelled.

### Docking the M^pro^ inhibitor 13b into the substrate binding cleft of M^pro^ validates the *AutoDock* docking protocol

The potential of HCV inhibitors to bind into the substrate-binding cleft of M^pro^ was assessed using *AutoDock* (Forli et al., 2016; Morris et al., 2009). To test the validity of our docking protocol, we carried out docking of a previously described inhibitor of the SARS-CoV-2 M^pro^, compound 13b (Supplementary Figure S1A), that also inhibits virus replication (Zhang et al., 2020a; Zhang et al., 2020b). Details of these control docking studies are provided in Supplementary Figures S1A – D, and in Supplementary Results. In the 1.75-Å X-ray crystal structure of the 13b-M^pro^ complex (Zhang et al., 2020b), approximately 50% of the 13b compound forms a reversible thiohemiketal covalent bond with the S^γ^ atom of catalytic residue Cys145 of the protease (Zhang et al., 2020b). The contribution of this partial covalent bond formation is difficult to assess, and our docking calculations determine only how well these drugs fit into the active site without the additional stabilization energy that may result from any covalent bond formation. The best-scoring docked conformation of 13b (Supplementary Figure S1C), has an *AutoDock* binding energy of -9.17 kcal/mol, and is similar but not identical to the pose observed in the X-ray crystal structure. Because the experimentally-determined pose is often not the one with the lowest docking energy, but rather is found among other highly-ranked poses (Kolb et al., 2009; Kolb and Irwin, 2009), we also examined other low energy poses. One such pose, which has a slightly higher binding energy of -9.03 kcal/mol, is almost identical to the pose observed in the crystal structure (Supplementary Figure S1D). Hence, binding poses very similar to that observed in the crystal structure are indeed included among the low energy poses generated by our *AutoDock* protocol.

### HCV protease inhibitors are predicted to bind snuggly into the substrate binding cleft of M^pro^

Having validated the use of *AutoDock* for docking of inhibitor 13b to M^pro^, and determining a representative *AutoDock* score for functionally-inhibitory binding (∼ - 9.0 kcal / mol) we carried out docking simulations with 10 HCV NS3/4A protease inhibitor drugs (Ozen et al., 2019). These 10 drugs have been approved for at least Phase 1 clinical trials, and some are FDA-approved prescription drugs used in treating HCV- infected patients (Table 1). The resulting docking scores are summarized in Figure 1D and Supplementary Table S1; and details of these docking poses are summarized in Supplementary Table S2. These results demonstrate that all of these HCV protease inhibitors have the potential to bind snuggly into the binding cleft of M^pro^, with extensive hydrogen-bonded and hydrophobic contacts, and predicted *AutoDock* energies of -8.37 to -11.01 kcal/mol, comparable to those obtained for the functional 13b inhibitor (∼ - 9.0 kcal/mol).

**Table 1.**
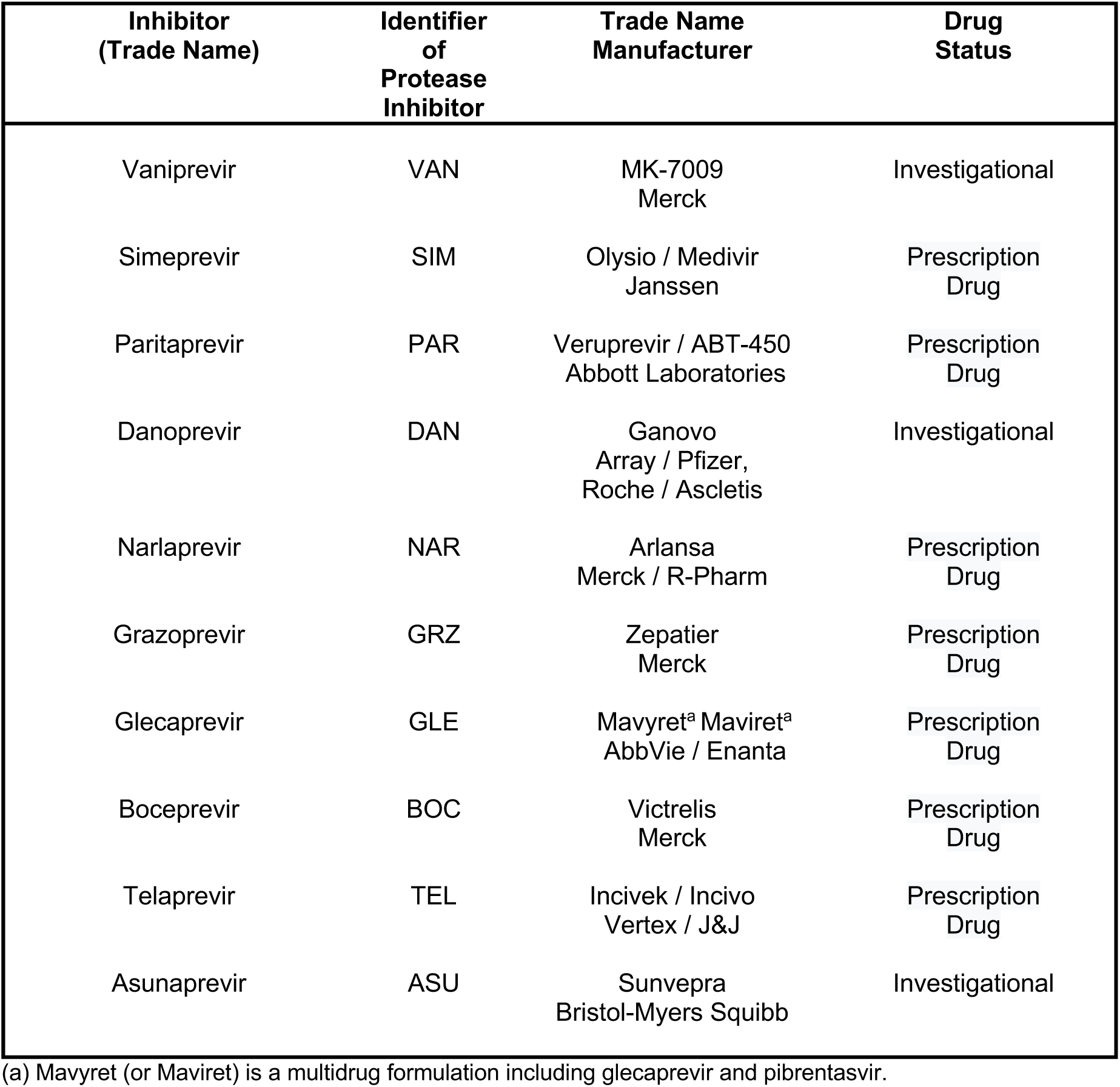
HCV 3C/4A Protease Inhibitors

| Inhibitor<br>(Trade Name) | Identifier<br>of<br>Protease<br>Inhibitor | Trade Name<br>Manufacturer | Drug<br>Status |
| --- | --- | --- | --- |
| Vaniprevir | VAN | MK-7009<br>Merck | Investigational |
| Simeprevir | SIM | Olysio / Medivir<br>Janssen | Prescription<br>Drug |
| Paritaprevir | PAR | Veruprevir / ABT-450<br>Abbott Laboratories | Prescription<br>Drug |
| Danoprevir | DAN | Ganovo<br>Array / Pfizer,<br>Roche / Ascletis | Investigational |
| Narlaprevir | NAR | Arlansa<br>Merck / R-Pharm | Prescription<br>Drug |
| Grazoprevir | GRZ | Zepatier<br>Merck | Prescription<br>Drug |
| Glecaprevir | GLE | Mavyret<sup>a</sup> Maviret<sup>a</sup><br>AbbVie / Enanta | Prescription<br>Drug |
| Boceprevir | BOC | Victrelis<br>Merck | Prescription<br>Drug |
| Telaprevir | TEL | Incivek / Incivo<br>Vertex / J&J | Prescription<br>Drug |
| Asunaprevir | ASU | Sunvepra<br>Bristol-Myers Squibb | Investigational |

2 (a) Mavyret (or Maviret) is a multidrug formulation including glecaprevir and pibrentasvir.

As we were preparing our work for publication, a 1.44 Å X-ray crystal structure of the complex of boceprevir bound to the SARS-CoV-2 M^pro^ was released in the Protein Data Bank [PDB id 6WNP (Anson et al., 2020; Anson and Mesecar, 2020)]. The boceprevir pose observed in this X-ray crystal structure is almost identical to the lowest energy pose (−9.03 kcal/mol) predicted by *AutoDock* (Figure 1E). In addition, both the docked and crystal structure binding poses for boceprevir near the active site of M^pro^ are very similar to its binding pose in the substrate binding cleft of HCV protease (Figure 1F), with an essentially identical hydrogen-bonding network between boceprevir and corresponding residues in each protease. Boceprevir is a alpha-ketoamide which potentially forms a covalent bond with the active site residue Cys145. Although this docking protocol does not include energetics or restraints for covalent bond formation, in the low energy pose of boceprevir bound to M^pro^, the alpha-keto amide carbon is positioned within 3.8 Å of the active site thiol sulfur atom. These blind tests of our docking results for boceprevir support the predictive value of docking results for the other HCV protease inhibitors.

Detailed images of the lowest-energy pose for seven of the ten complexes are shown in Supplementary Figure S2, and three representative complexes, for vaniprevir, simeprevir, and narlaprevir are illustrated in Figure 1G - I. In addition, a summary of *AutoDock* scores (Supplementary Table S1), key intermolecular contacts in each of the ten complexes (Supplementary Table S2), and descriptions of these binding poses, are presented in Supplementary Information. In this analysis, we paid particular attention to key details of the lowest energy pose for each inhibitor, including interactions with the sidechains of catalytic dyad residues His41 and Cys145, and hydrogen-bonded interactions with the backbone amides of Gly143, Ser144 and Cys145, which form the oxyanion hole of this cysteine protease (Zhang et al., 2020b). All these HCV protease inhibitors form extensive hydrogen-bonded networks and hydrophobic interactions within the substrate binding cleft, and would occlude access of polypeptide substrates to the active-site residues His41 and Cys145 (as shown for example in Figure 1G-I). While the three HCV protease inhibitors with alpha-keto amide functional groups, namely, boceprevir, narlaprevir, and telaprevir, have lowest-energy or low-energy poses that could form covalent bonds with the active-site Cys145 thiol, the seven other HCV protease inhibitors are non-covalent inhibitors that would not form such thioester bonds. From these virtual docking studies, we conclude that all ten of these HCV protease inhibitors have the potential to bind snuggly into the substrate binding cleft of M^pro^, and to inhibit binding of its substrates.

### Seven HCV drugs inhibit M^pro^ protease activity

Inhibition activity of HCV drugs against M^pro^ was initially assessed using a protease assay based on Föster resonance energy transfer (FRET) using the peptide substrate Dabsyl-KTSAVLQ/SGFRKME-(Edans), containing a canonical M^pro^ protease recognition site. Rates of hydrolysis were determined for 90 minutes, and initial velocities were used to assess the relative inhibitory activities of the ten HCV protease inhibitors. The percent M^pro^ activity denotes the ratio of initial hydrolysis velocities in the presence of 20 μM drug to the velocities in the presence of only the DMSO solvent. These assays used 10 nM M^pro^ and 20 μM peptide substrate, at a final DMSO concentration of 1.0%. Under these conditions, HCV protease inhibitors narlaprevir, boceprevir, and telaprevir have significant enzyme inhibition activity, with IC_50_ values of 2.2 ± 0.4 μM, 2.9 ± 0.6 μM, and 18.7 ± 6.4 μM, respectively (Figures 2A, B). In contrast, little or no inhibition activity was detected with the other seven HCV protease inhibitors (Figure 2A and Supplementary Figure S3A).

**Fig. 2.**
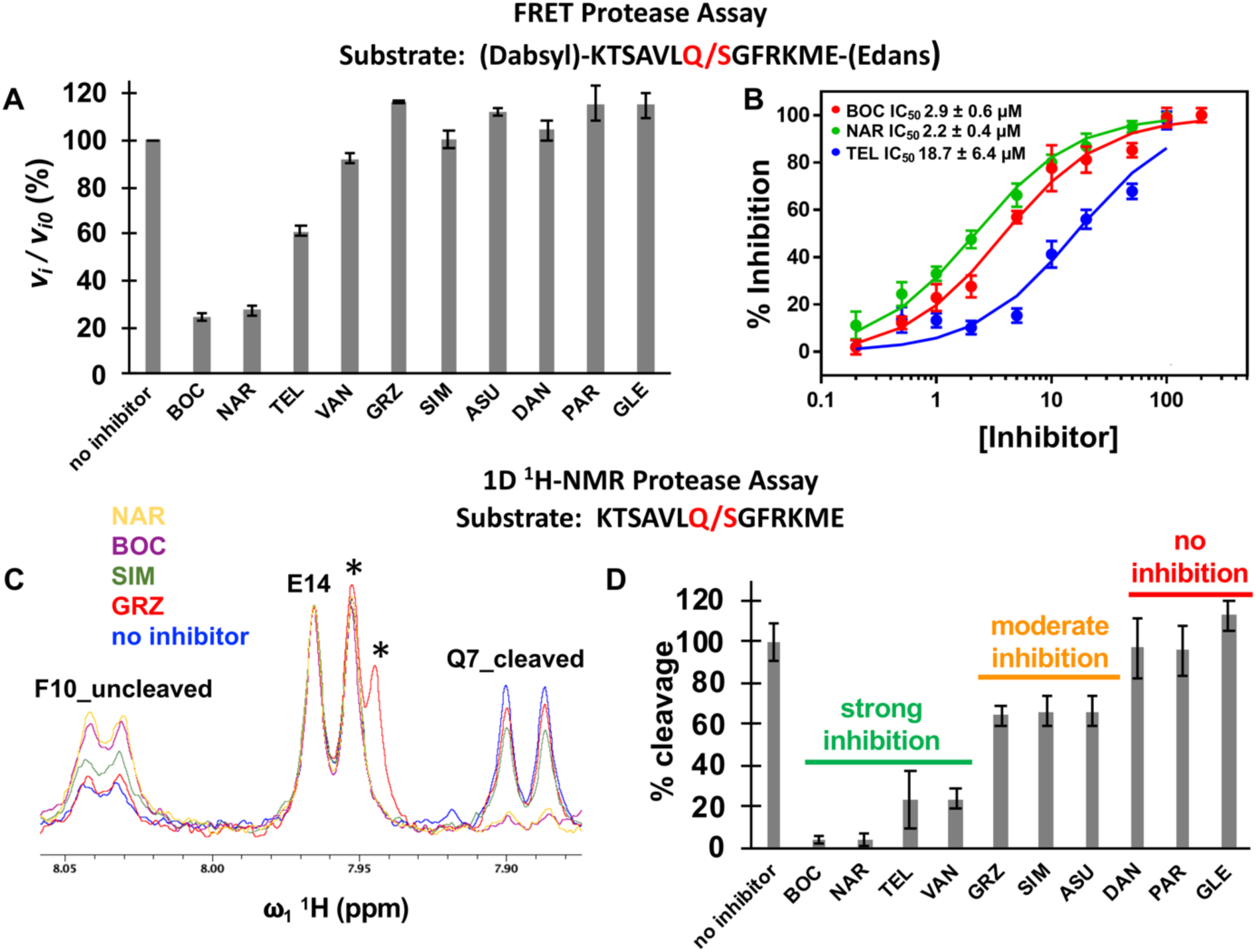
HCV protease inhibitors inhibit SARS-CoV-2 M^pro^. (A) Initial rates of proteolysis of a peptide substrate by M^pro^ in the presence of 20 mM inhibitor concentrations (v_i_) relative to initial rate in the absence of inhibitor (v_i,o_), at 25° C. (B) Dose response curves based on FRET assay for inhibition of M^pro^ using the indicated peptide substrate, by narlaprevir (NAR), boceprevir (BOC), and telaprevir (TEL). In these FRET assays, the M^pro^ concentration is 10 nM, and the substrate concentration is 20 mM. (C) 1D ^1^H-NMR assay for hydrolysis of the indicated peptide substrate. The amide proton doublets of Phe-10 prior to cleavage (F10_uncleaved) or of Gln-7 after cleavage (Q7_cleaved) provide well-resolved resonances for monitoring the proteolysis reaction. The amide proton doublet of Glu-14 (E14) is not perturbed by cleavage, and provides an internal intensity calibration control. Grazoprevir (GRZ) has resonances (labeled by *) that overlap with the upfield component of the E14 doublet. (D) Percent cleavage of the indicated peptide substrate by M^pro^ after 30 min at 25° C. These NMR studies used an M^pro^ enzyme concentration of 100 nM, peptide substrate concentration of 50 mM, and inhibitor drug concentrations of 50 mM.

While very sensitive, this FRET assay is complicated by the fact that, aside from narlaprevir, boceprevir, and telaprevir, the other HCV drugs have significant intrinsic fluorescence that interferes with FRET measurements using this peptide substrate (compare fluorescence intensity at time t = 0 sec in Supplementary Figure S3A). Consequently, M^pro^ inhibition by some of these drugs might be masked or interfered with by fluorophore interactions. To overcome this, we also developed a 1D ^1^H-NMR assay, using the peptide substrate KTSAVLQ/SGFRKME that lacks the Dabsyl and Edans N-terminal and C-terminal tags, as outlined in Figure 2C and Supplementary Figure S4. In this 1D ^1^H-NMR assay, M^pro^ activity was defined as the ratio of hydrolyzed peptide observed after 30 min in the presence of drug, to the amount of hydrolyzed peptide observed when the enzyme is incubated with DMSO alone. These assays used 100 nM M^pro^, 50 μM peptide substrate, and 50 μM drug (or DMSO solvent alone), at a final DMSO concentration of 1.0%. Under these conditions, narlaprevir, boceprevir, and telaprevir have substantial enzyme inhibition activity, as was the case in the FRET assay. In addition, in the NMR assay substantial inhibitory activity was also observed for vaniprevir, and moderate inhibitory activity was observed for grazoprevir, simeprevir, and asunaprevir (Figure 2D). This moderate inhibition was not detected in the FRET assay. Danoprevir, paritaprevir, and glecaprevir had little or no detectable M^pro^ inhibitory activity. From these studies we conclude that seven HCV drugs (*viz* boceprevir, narlaprevir, telaprevir, vaniprevir, grazoprevir, simeprevir, and asunaprevir) inhibit SARS-CoV-2 M^pro^ strongly or moderately under the conditions tested.

### Three HCV protease inhibitors inhibit not only M^pro^ but also PL^pro^

Although the active site of PL^pro^ does not share structural similarity with the HCV protease, it is possible that some HCV protease inhibitors can also bind into the active site of PL^pro^. Accordingly, we carried out virtual docking studies of these same ten HCV drugs into the substrate binding cleft of PL^pro^, using protocols similar to those developed in virtual docking studies with M^pro^. The PL^pro^ inhibitor GRL-0617, for which a crystal structure bound to PL^pro^ is available in the PDB (Fu and Huang, 2020), was used to assess the docking protocol and to determine a reference *AutoDock* score of -7.54 kcal / mol. The scores of docking poses for HCV drugs, summarized in Figure 3A and Supplementary Table S1, range from – 5.56 kcal / mole for boceprevir and narlaprevir, to much more favorable values of < – 8 kcal / mol for others including vaniprevir, grazoprevir, simeprevir, and paritaprevir. The most favorable docking poses for simeprevir and vaniprevir are shown in Figures 3B - C, and for the remaining complexes in Supplementary Figure S5. These results indicate that, surprisingly, some HCV protease inhibitors may bind in the active sites of both M^pro^ and PL^pro^.

**Fig 3.**
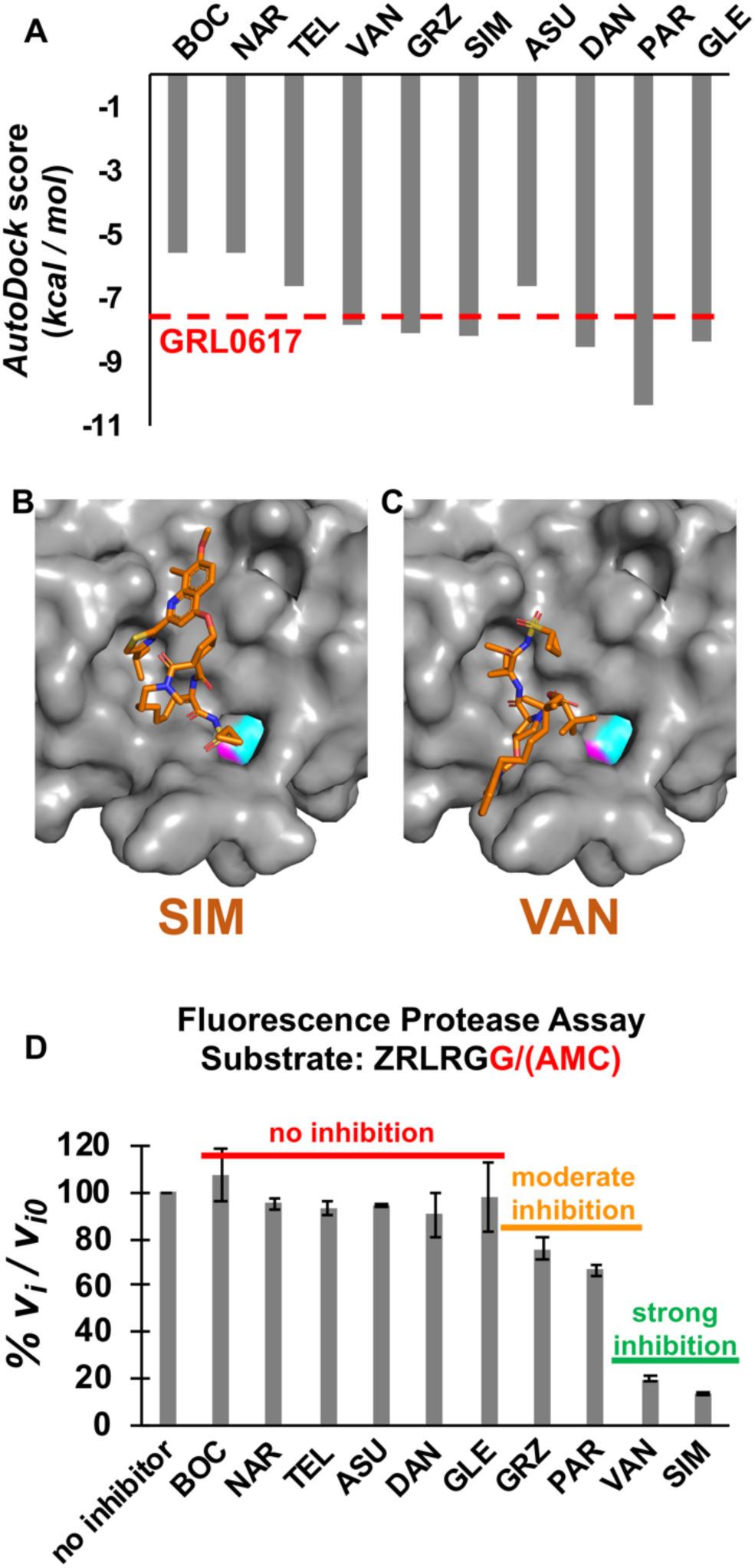
HCV protease inhibitors also inhibit SARS-CoV-2 PL^pro^. (A) *AutoDock* docking scores for 10 HCV protease inhibitors in the substrate binding cleft of SARS-CoV-2 PL^pro^. The docking score for PL^pro^ inhibitor GRL-0617 (*AutoDock score* = -7.54 kcal / mol) is also shown as a horizontal red dashed line. (B) Best-scoring *AutoDock* poses for complexes of simeprevir (SIM) and vaniprevir (VAN) with PL^pro^. (C) Initial rates of proteolysis of a peptide substrate by PL^pro^ in the presence of 20 µM inhibitor concentrations (*v*_i_) relative to initial rate in the absence of inhibitor (*v*_i,o_), at 25° C. Grazoprevir inhibition of PL^pro^ was even stronger (*v*_o_/*v*_o,i_ = 32 ± 7%) at 100 µM drug concentration (Supplementary Figure S3B).

Based on these docking results, we anticipated that several HCV protease inhibitors, not including bocepriver or narlaprevir, might inhibit PL^pro^ protease activity. To test this hypothesis, fluorescence assays of PL^pro^ inhibition were carried out, using the substrate ZRLRGG/AMC (Z - carboxybenzyl; AMC - 7-Amino-4-methylcoumarin) (Anson et al., 2020) containing a natural/canonical PL^pro^ protease recognition site. In this assay, the AMC group is quenched when covalently attached to the peptide, and increases fluorescence upon proteolytic cleavage. Of the HCV drugs tested, four drugs predicted to bind well into the active site of PL^pro^, vaniprevir, simeprevir, paritaprevir, and grazoprevir, inhibit PL^pro^ protease activity (Figure 3C and Supplementary Figure S3B). Hence, vaniprevir, simeprevir, and grazoprevir inhibited both M^pro^ and PL^pro^, while paritaprevir inhibited PL^pro^, but not M^pro^. Vaniprevir and simeprevir are more effective PL^pro^ inhibitors than grazoprevir or paritaprevir under these assay conditions.

### Seven HCV drugs inhibit SARS-CoV-2 replication in Vero and/or human 293T cells

The motivation for the docking and biophysical studies described above was to identify HCV drugs with the potential to inhibit SARS-CoV-2 virus replication. For antiviral assays, Vero E6 cells, or human 293T cells expressing the SARS-CoV-2 ACE2 receptor, were grown in 96-well plates, and were incubated with various levels of a HCV protease inhibitor for 2 hours. Cells were then infected with SARS-CoV-2 virus at the indicated multiplicity of infection [moi, plaque-forming units (pfu)/cell] and incubated for the indicated times at 37^0^ C in the presence of the inhibitor. Virus-infected cells were then identified by immunofluorescence using an antibody specific for the viral nucleoprotein. Inhibition of viral replication was quantified by determining the percentage of positive infected cells at the end of the incubation period in the presence of the compound, as compared with the number of infected cells in its absence. To determine whether a HCV drug was cytotoxic, uninfected Vero E6 or human 293T cells were incubated with the same levels of the compounds for the same length of time, and cytotoxicity was measured using an MTT assay (Roche). In all of these replication assays, remdesivir was used as a positive control. Further details of these replication assays are provided in STAR Methods.

Viral replication inhibition data in Vero E6 cells are presented in Figure 4. Three of the HCV protease inhibitors tested, grazoprevir, asunaprevir, and boceprevir, inhibited SARS-CoV-2 virus replication at concentrations lower than the concentrations that cause significant cytotoxicity. Two other HCV protease inhibitors, simeprevir and vaniprevir, inhibited virus replication with even lower IC_50_ values, but some cytotoxicity was also observed. These five HCV drugs have IC_50_ values for inhibiting SARS-CoV-2 replication of 4.2 to 19.6 μM. The remaining three HCV drugs tested, telaprevir, glecaprevir, and danoprevir, did not inhibit virus replication in Vero E6 cells, even at a maximum drug concentrations of 50 μM.

**Fig. 4.**
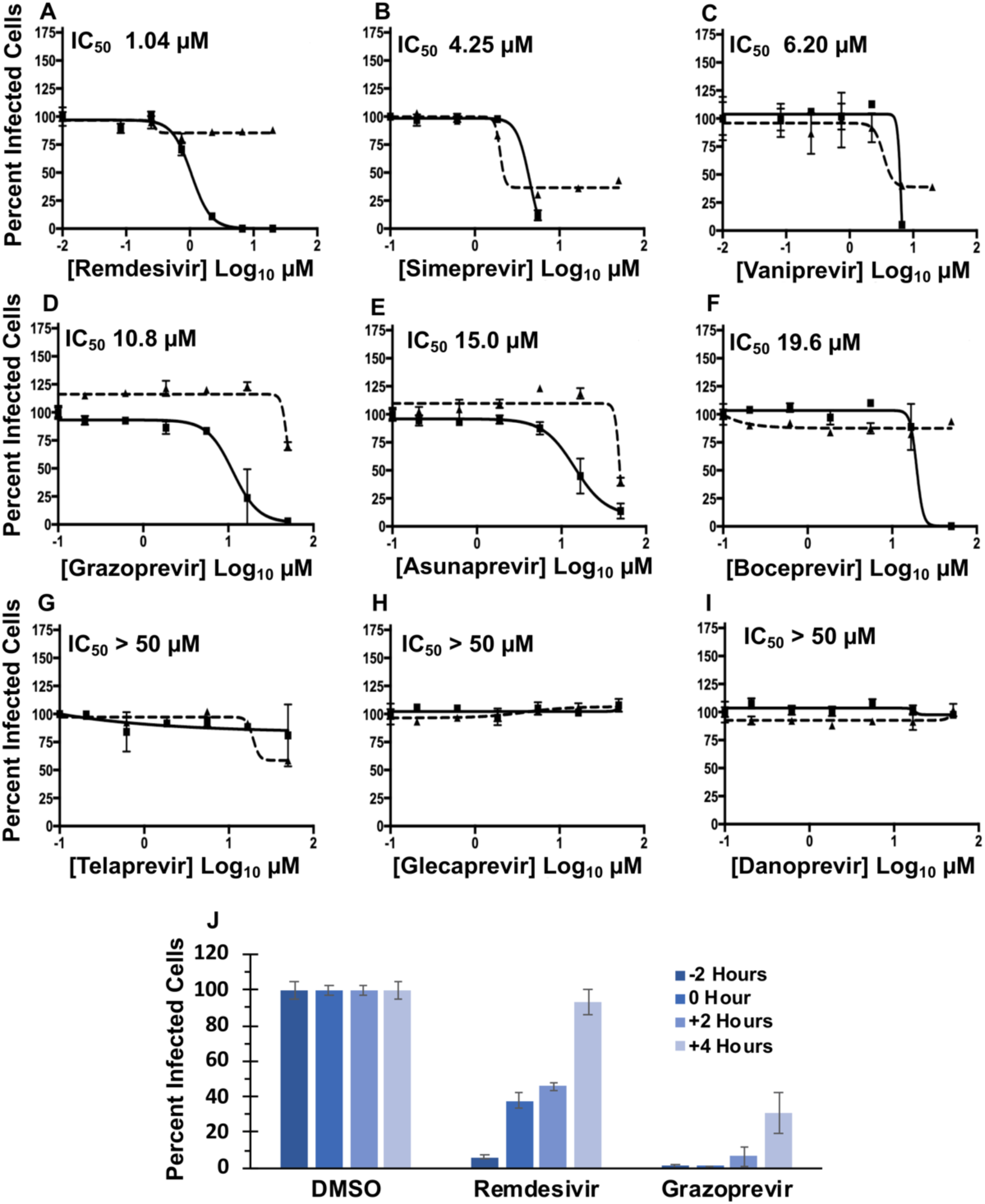
Antiviral activity of HCV protease inhibitors in Vero E6 cells. (A – I) Inhibition of viral replication was determined in a concentration-dependent matter in Vero E6 cells. Replication assays were performed at multiplicity of infection (moi) of 0.025 plaque forming units (pfus) /cell. In all panels, viral infectivity is shown as a solid line and cell viability as a dashed line. Data = mean ± SD; n = 3 independent samples. The estimated IC_50_ is labeled in the upper left corner of each plot. Remdesivir is included as a standard of care control. (J) Time of addition assay. 10 mM of remdesivir or 50 mM of grazoprevir were added to cells at the indicted time points before (- 2 hrs), at (0 hrs), or after (+2 or +4 hrs) viral infection.

We next determined whether the inhibition of SARS-CoV-2 replication by a representative HCV drug occurs at steps after virus entry, as would be expected for inhibitors of a viral protease that is produced only after infection. Accordingly, we performed a time of addition assay using grazoprevir as the inhibitor. In a single cycle (moi of 2 pfu/cell) infection, grazoprevir was added 2 hours prior to infection, at the time of infection, or at 2 or 4 hours post infection. Grazoprevir inhibited virus replication at all time points tested (Figure 4J). Because virus replication was inhibited by the addition of grazoprevir as late as 4 hours post infection, it is unlikely that grazoprevir is inhibiting viral entry, which should be completed by this time post-infection. These results indicate that grazoprevir inhibits steps during the replication stage of the viral life cycle, hence implicating virus-encoded proteases in the inhibition of virus replication.

We also determined whether the HCV drugs exhibited similar antiviral activities in human cells, specifically human 293T cells expressing the ACE2 receptor (Figure 5). We included in these antiviral assays another HCV protease inhibitor, narlaprevir, which we observed to be a strong M^pro^ inhibitor (Figure 2B). Again, simeprevir and vaniprevir were the most effective inhibitors of virus replication, with IC_50_ values of 2.3 and 3.0 μM, respectively, and with considerably reduced cytotoxicity as compared to that in Vero cells. Boceprevir and narleprevir, which are strong M^pro^ inhibitors (Figure 2A,B), were less effective inhibitors of virus replication, with IC_50_s of 5.4 and 15 μM. The other reversible covalent M^pro^ inhibitor, telaprevir, had an even lower IC_50_ of 20.5 μM. Grazoprevir had considerably lower antiviral activity in human 293T cells than in Vero cells (compare Figures 4D and 5F). Thus, six HCV drugs inhibited SARS-CoV-2 replication in human 293T cells, with IC_50_ values ranging from 2.3 to 20.5 μM. Glecaprevir and danoprevir did not inhibit virus replication in 293T cells, as was also the case in Vero cells.

**Fig. 5.**
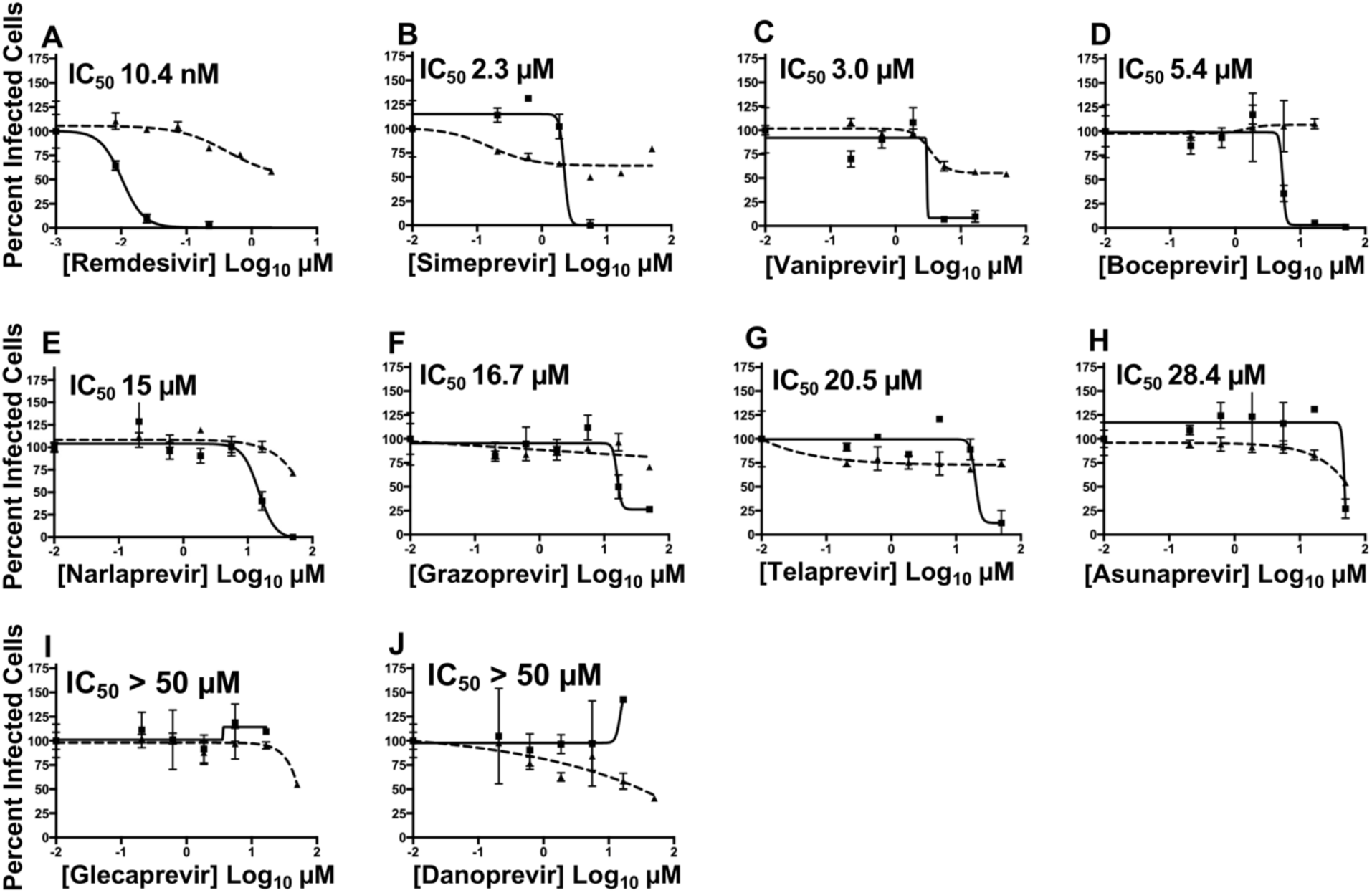
Antiviral activity of HCV protease inhibitors in HEK293T cells. (A – J) Inhibition of viral replication was determined in a concentration-dependent matter in HEK293T cells. Replication assays were performed at multiplicity of infection (moi) of 0.025 plaque forming units (pfus) /cell. In all panels, viral infectivity is shown as a solid line and cell viability as a dashed line. Data = mean ± SD; n = 3 independent samples. Calculated IC_50_ is indicated in the upper left corner of each plot. Remdesivir is included as a standard of care control.

### Simeprevir, and grazoprevir synergize with remdesivir to increase inhibition of SARS-CoV-2 virus replication

Because M^pro^ and PL^pro^ generate either the RNA polymerase itself, or the proteins that constitute the replication organelles required for polymerase function, we predicted that HCV drugs that inhibit one or both of these viral proteases might be synergistic with inhibitors of the viral polymerase like remdesivir. To test this hypothesis, we carried out antiviral combination assays in Vero cells of simeprevir, grazoprevir, or boceprevir with remdesivir (Figure 6). To assess synergy, two analyses are required. In one analysis the IC_90_ of the remdesivir was measured in the presence of increasing concentrations of each of these HCV drugs (Figure 6A-C). These results demonstrate that simeprevir or grazoprevir increase the antiviral activity of remdesivir. For example, in the presence of 1.25 µM simeprevir, approximately 10-fold less remdesivir is required for the same antiviral effect achieved in the absence of simeprevir (Figure 6A). Surprisingly, although boceprevir is a much better inhibitor of M^pro^ than either simeprevir or grazoprevir, boceprevir did not significantly affect the antiviral activity of remdesivir (Figure 6C). In the second analysis, the IC_90_ concentration of each HCV drug was determined in the presence of increasing concentrations of remdesivir (Figures 6D-F). Remdesivir increased the antiviral activity of simeprevir and grazoprevir. For example addition of 1.25 µM remdesivir substantially reduces the concentration of simeprevir needed to achieve IC_90_ conditions (Figure 6D). In contrast, remdesivir did not significantly affect the antiviral activity of boceprevir (Figure 6F).

**Fig. 6.**
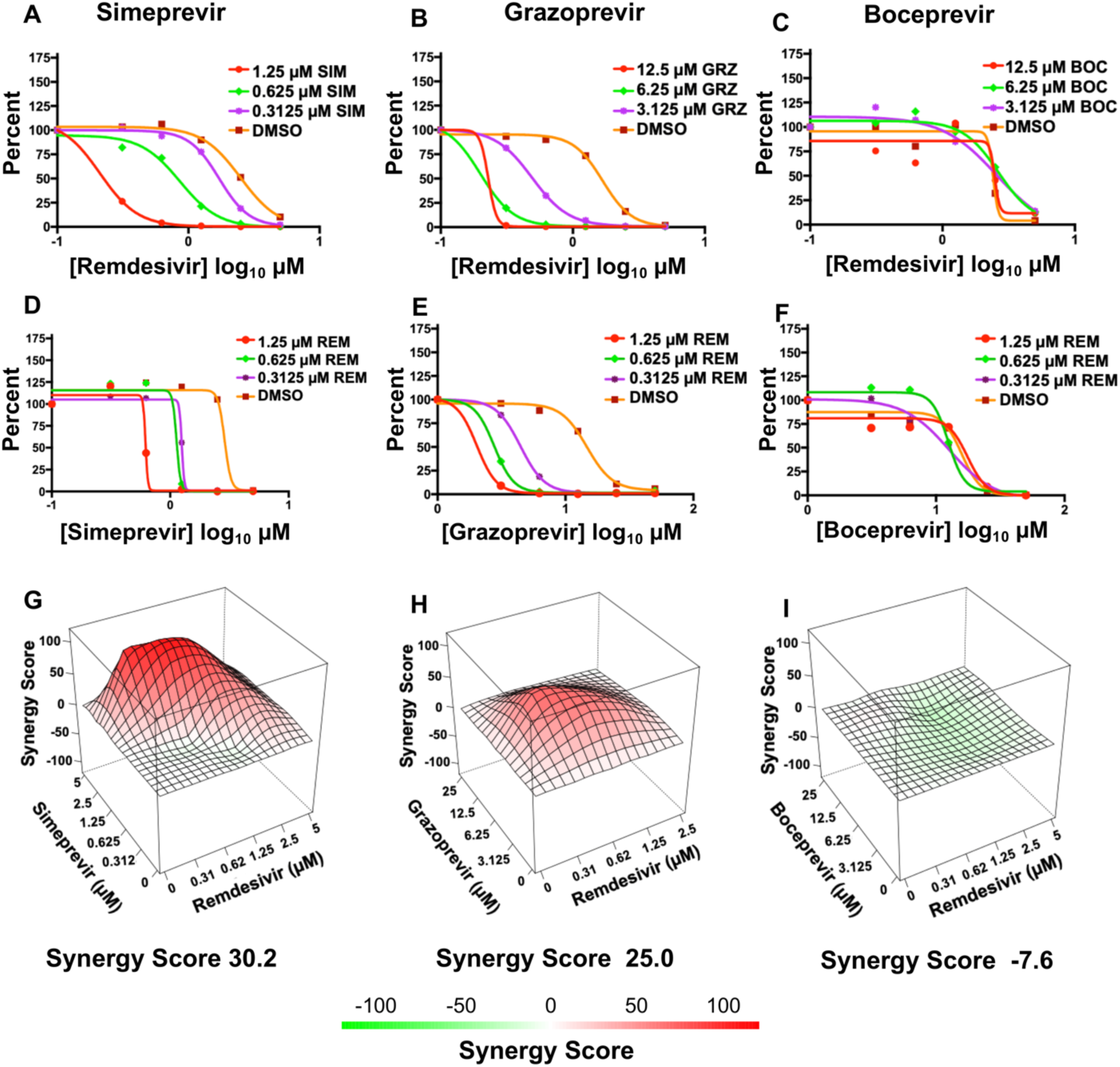
Simeprevir and grazoprevir are synergistic with remdesivir in Vero E6 cells. (A - C) SARS-CoV-2 inhibition by remdesivir in the presence of increasing concentrations of simeprevir, grazoprevir, or boceprevir. (D - F) SARS-CoV-2 inhibition by simeprevir, grazoprevir, or boceprevir in the presence of increasing concentrations of remdesivir. (G - I) Synergy landscapes and combination scores generated by the ZIP method using the program *SynergyFinder* (Ianevski et al., 2020). Red surfaces denote a synergistic interaction, and green surfaces an antagonistic interaction. In this model a synergistic interaction between drugs has a score greater than +10; an additive interaction has a score between -10 to +10; and an antagonistic interaction has a score of less than -10 (Ianevski et al., 2020).

These combination antiviral assays indicate that simeprevir and grazoprevir, but not boceprevir, act synergistically with remdesivir to inhibit virus replication. As confirmation, we subjected these results to analysis by the Zero Interaction Potency (ZIP) model for synergy (Ianevski et al., 2020). In the landscapes generated by this model that are shown in Figure 6G - I, red denotes a synergistic interaction, and green denotes an antagonistic interaction. In this model a synergistic interaction between drugs has a score greater than +10; an additive interaction has a score between -10 to +10; and an antagonistic interaction has a score of less than -10. The landscapes for the interaction of remdesivir with both simeprevir and grazoprevir are red, with synergy scores of +30.2 and +25.0, respectively, denoting moderate synergism. In contrast, although the landscape for the interaction of remdesivir with boceprevir suggests weak antagonism (Figure 6I), the score of the interaction is -7.6, which we interpret as an additive interaction.

We also carried out combination antiviral assays in human 293T cells. The interaction between remdesivir and grazoprevir in inhibiting virus replication in the human cells was also synergistic, with a red landscape and a synergy score of +20.3 (Supplementary Figure S6A). The interaction between remdesivir and vaniprevir in inhibiting virus replication was also analyzed in 293T cells; it was weakly synergistic, with a score of +10.9. Vaniprevir has an additive effect or, at best, very weak synergy with remdesivir in human 293T cells. Consequently, two HCV drugs, simeprevir and grazoprevir, act synergistically with remdesivir to inhibit SARS-CoV-2 virus replication in Vero and/or human 293T cells.

## Discussion

To provide antiviral drugs that can be rapidly deployed to combat the COVID-19 pandemic, we carried out the present study to identify currently available drugs that could potentially be repurposed as inhibitors of the pandemic SARS-CoV-2 virus that causes COVID-19 disease. Instead of screening libraries of current drugs, we took a different approach to identify candidate drugs. Specifically, we initiated our search based on the striking similarity of the substrate binding clefts of the SARS-CoV-2 M^pro^ and HCV proteases. The HCV protease is a serine-protease, with catalytic triad His57, Asp81, and Ser139, while the SARS-CoV-2 M^pro^ is a cysteine protease, with catalytic dyad residues His41, Cys145 (Jin et al., 2020; Zhang et al., 2020b), While the two structures have similar structural architectures (i.e. organization of secondary structure elements) (Orengo et al., 1997), their secondary structure topologies (i.e. how the secondary structure elements are linked together) are very different (Supplementary Figure S7A), and do not indicate a divergent evolutionary relationship. However, a structure-based sequence alignment, generated from these superimposed three- dimensional structures, aligns catalytic residues His41 / His57 and Cys145 / Ser139 (Supplementary Figure S7B), where each aligned pair is found in a similar secondary structure element. Hence, it appears that the similarities in the overall structures and relative orientations of domains I and II, the features of the substrate binding cleft located between these domains, and the relative positions of catalytic residues in these enzymes, is a result of convergent evolution (Kester and Matthews, 1977; Robertus et al., 1972) to create a similar protease active site.

As predicted by the striking similarity of the substrate binding clefts of the SARS- CoV-2 M^pro^ and HCV proteases, our virtual docking experiments showed that ten HCV protease inhibitors can be docked snuggly into the substrate binding cleft of M^pro^, and hence would have the potential to inhibit binding of the M^pro^ substrate, thereby inhibiting proteolytic cleavage of the substrate. In fact, we showed that four HCV drugs, boceprevir, narlaprevir, telaprevir, and vaniprevir strongly inhibit SARS-CoV-2 M^pro^ protease activity, and that three other HCV drugs, grazoprevir, simeprevir and asunaprevir moderately inhibit M^pro^ activity. Boceprevir, narleprevir and telaprevir are *α*- keto amides, which can form a reversible covalent bond with the active site Cys thiol of M^pro^. The other four HCV drugs that inhibit the M^pro^ protease cannot form a covalent bond active site Cys thiol of M^pro^. These seven HCV drugs inhibit virus replication in Vero and/or human 293T cells expressing the ACE2 receptor. Although the active site of PL^pro^ does not share structural similarity with the HCV NS3/4A protease, four HCV drugs, simeprevir, grazoprevir, vaniprevir, and paritaprevir can be docked with energetically-favorable scores into the active site of PL^pro^, and inhibit PL^pro^ protease activity *in vitro*. None of these four inhibitors can form covalent bonds with the active- site Cys residue of PL^pro^. Vaniprevir is the only one of the tested HCV drugs that strongly inhibits both M^pro^ and PL^pro^, presumably accounting for its strong inhibition of virus replication.

Other groups have also recently reported that some of these same HCV protease inhibitors can inhibit M^pro^ or PL^pro^ protease activities, and that boceprevir can inhibit SARS-CoV-2 replication (Anson et al., 2020; Fu et al., 2020; Ma et al., 2020). While all of these studies report boceprevir as a moderately-potent inhibitor of M^pro^, there are inconsistent reports of the effectiveness of some of the other HCV protease inhibitors reported here as inhibitors of M^pro^ and/or PL^pro^. These inconsistencies likely arise from details of the different assays that have been used. For example, our ^1^H-NMR assay could identify moderate inhibition of M^pro^ by simeprevir, grazoprevir, and asunaprevir HCV drugs that was not detected by our FRET assay. This is not surprising, since both the FRET and NMR assays are competition assays, which will give different results for substrates that have different binding affinities. The ability of HCV drugs to inhibit M^pro^ may also depend on other details of the assay conditions, most notably the enzyme, substrate, and drug concentrations which may affect the monomer-dimer equilibrium of M^pro^ (Grum-Tokars et al., 2008).

PL^pro^ not only cleaves viral polyproteins to generate crucial viral proteins, but also cleaves the ubiquitin-like ISG15 protein from viral proteins (Daczkowski et al., 2017; Lindner et al., 2007). ISG15 conjugation in infected cells results in a dominant-negative effect on the functions of viral proteins (Zhao et al., 2016), and removal of ISG15 from viral proteins by PL^pro^ would restore these functions of these viral proteins. In addition, the resulting free ISG15 would be secreted from infected cells and would bind to the LFA-1 receptor on immune cells, causing the release of interferon-*γ* and inflammatory cytokines (Swaim et al., 2020; Swaim et al., 2017). The release of these cytokines could contribute to the strong inflammatory response, the so-called cytokine storm, that has been implicated in the severity of COVID-19 disease (McGonagle et al., 2020). Inhibition of PL^pro^ by a HCV drug should also inhibit the release of interferon-*γ* and inflammatory cytokines, potentially mitigating the cytokine storm.

HCV drugs that inhibit M^pro^ and/or PL^pro^ would be expected to inhibit viral RNA polymerase function in infected cells by inhibiting the generation of protein subunits of the viral polymerase and/or by inhibiting the generation of the integral-membrane non- structural viral proteins that form the replication organelles (double membrane vesicles) required for polymerase function. Because of the resulting reduction of functional viral polymerase, the amount of remdesivir needed for inhibition of virus replication would likely be reduced. Viral replication assays using combinations of drugs allowed us to assess if the interactions between HCV drugs and remdesivir are additive or synergistic, i.e. resulting in an inhibition of virus replication that is greater than the sum of the inhibitions caused by the HCV drug and remdesivir. Surprisingly, we found that these inhibitory effects are additive or synergistic depending on which HCV drug is used to inhibit virus replication. Boceprevir, which strongly inhibits only M^pro^, and vaniprevir, which strongly inhibits both M^pro^ and PL^pro^, act additively with remdesivir to inhibit virus replication. In contrast, simeprevir and grazoprevir, which moderately inhibit M^pro^ and either strongly or moderately inhibit PL^pro^, act synergistically with remdesivir to inhibit virus replication. The basis for the different interactions between remdesivir and particular HCV drugs, and the mechanism(s) of synergy, needs to be further explored.

The HCV drugs that are strongly synergistic with remdesivir are most pertinent for the goal of the present study to identify available drugs that can be repurposed as SARS-CoV-2 antivirals and/or as lead molecules for new drug development. Repurposed drugs like the HCV drugs in the present study may not have sufficient inhibitory activity on their own to achieve clinical efficacy. Synergy with remdesivir increases the potency of both the repurposed HCV drug and remdesivir. We identified two HCV drugs, simeprevir and grazoprevir, that act synergistically with remdesivir to inhibit SARS-CoV-2 virus replication. Of these two, simeprevir may be the better choice as a repurposed drug because it effectively inhibits SARS-CoV-2 virus replication in human cells at much lower concentrations than grazoprevir. Consequently, the combination of simeprevir and remdesivir could potentially function as an antiviral against SARS-CoV-2 while more specific SARS-CoV-2 antivirals are being developed. Simeprevir, which is an oral drug, might also be combined with an oral polymerase inhibitor rather than with remdesivir, which has to be administered intravenously. One such oral polymerase inhibitor, molnupiravir (EIDD-2801) (Sheahan et al., 2020), which is currently in late stage clinical trials, could potentially be combined with for clinical applications. Accordingly, a combination of simeprevir and molnupiravir, both of which have oral bioavailability, could be assessed for outpatient use.

## Acknowledgements

Drs. G. Chalmers, Y.P. Huang, G. Liu, L. Ma, C. Sander, S. Shukla, G.V.T Swapna, and R. Xiao for helpful discussions, suggestions, and comments on the manuscript. We also thank J. Hunt, S. Krishna, D. Aalberts, Greg Boël for a generous gift of SARS-CoV2 PL^pro^ enzyme, and R. Albert for support with the BSL3 facility and procedures at the Icahn School of Medicine at Mount Sinai, New York. This research was supported by grants from the National Institutes of Health (R01-GM120574 to GTM) and RPI Center for Computational Innovations (to KB and GTM). This research was also partly funded by CRIP (Center for Research for Influenza Pathogenesis), a NIAID supported Center of Excellence for Influenza Research and Surveillance (CEIRS, contract # HHSN272201400008C), by DARPA grant HR0011-19-2-0020, by supplements to NIAID grant U19AI142733 U19AI135972 and DoD grant W81XWH-20-1-0270, and by the generous support of the JPB Foundation, the Open Philanthropy Project (research grant 2020-215611 (5384)), and anonymous donors to AG-S.

## Author Contributions (Categories are defined by Cell Journal)

Conceptualization, KB, KW, BH, CAR, AG-S, RMK, GTM; Methodology, KB, KW, BH, TBA, TR, RR, CAR, AG-S, RMK, GTM; Validation, KB, KW, BH, CAR, AG-S, RMK, GTM; Formal Analysis, KB, KW, BH, TR, CAR, AG-S, RMK, GTM; Investigation, KB, KW, BH, RR, TR, EM, TK, LM; Resources, CAR, AG-S, and GTM; Writing-original draft, KB, KW, AG-S, RMK, GTM; Writing, Review and Editing: KB, KW, BH, CAR, AG-S, RMK, GTM; Visualization, KB, KW, BH, CAR, AG-S, RMK, GTM; Supervision, KW, CAR, AG-S, GTM; Funding Acquisition, CAR, AG-S, GTM.

## STAR METHODS

**Table SM1 - Key Resources**

3 Table SM1 - Key Resources
| REAGENT or RESOURCE | SOURCE | IDENTIFIER |
| --- | --- | --- |
| **Cell Lines** |  |  |
| *E. coli* BL21(DE3) | New England Biolabs Inc | C2527 |
| Vero E6 cells | ATCC | CRL-1586 |
| HEK293T cells | ATCC | CRL-3216 |
| **Chemicals** |  |  |
| Luria Broth (LB) | Fischer BioReagents | BP1426-2 |
| LB Agar | VWR Amresco | J104 |
| Ampicillin | Research Products International | A40040 |
| IPTG | Research Products International | 156000 |
| Sodium chloride | VWR Amresco | X190 |
| Tris | MP Biomedicals LLC | 819263 |
| TCEP | VWR Amresco | K831 |
| Imidazole | Alfa Aesar | A10221 |
| DTT | VWR Life Sciences | 0281 |
| EDTA | VWR Amresco | M101 |
| HEPES | Chem Impex Int'l Inc | 00174 |
| Bovine Serum Albumin | VWR Amresco | 0322 |
| D<sub>2</sub>O | Sigma-Aldrich | 343773 |
| d<sub>6</sub>DMSO | Cambridge Isotope Laboratories Inc. | DLM-10 |
| Ni-NTA resin | MCLAB | NiNTA-100 |
| Sodium Azide | VWR Amresco | 0639 |
| NaOH | VWR Life Sciences | M137 |
| **SDS PAGE gel and staining** |  |  |
| Acrylamide/bis-acrylamide | VWR Life Sciences | 0311 |
| Sodium Dodecyl Sulphate | VWR Life Sciences | 0837 |
| Ammonium persulphate (APS) | VWR Amresco | M133 |
| TEMED | Thermo Scientific | 17919 |
| Bromophenol blue | VWR Amresco | M131 |
| Glycerol | Fisher Chemical | G33 |
| Glycine | VWR Life Sciences | 0176 |
| Glacial acetic acid | Fisher Chemical | A38-212 |
| Isopropanol | Fisher Bioreagents | BP2618 |
| HCl | MP Biomedicals LLC | 194054 |
| **Inhibitors** |  |  |
| Asunaprevir | Selleckchem Inc. | S4935 |
| Boceprevir | Selleckchem Inc. | S3733 |
| Danoprevir | Selleckchem Inc. | S1183 |
| Glecaprevir | Selleckchem Inc. | S5720 |
| Grazoprevir | Selleckchem Inc. | S3728 |
| Paritaprevir | Selleckchem Inc. | S5404 |
| Simeprevir | Selleckchem Inc. | S5015 |
| Telaprevir | Selleckchem Inc. | S1538 |
| Narlaprevir | MedChemExpress | HY-10300 |
| Vaniprevir | Addoq | A11600-5 |
| Remdesivir | Medkoo Bioscience Inc. | 329511 |
| **Plasmids** |  |  |
| M<sup>pro</sup> expression vector<br>GTM_COV2_NSP5_001 | GenScript Inc. | Custom synthesis |
| **Peptide substrates** |  |  |
| KTSAVLQ/SGFRKME | GenScript Inc. | Custom synthesis |
| Dabcyl-KTSAVLQ/SGFRKME-Edans | GenScript Inc. | Custom synthesis |
| zRLRGG/AMC | Bachem Americas Inc. | 4027158.0025 |
| **Software and Algorithms** |  |  |
| Topspin 3.2.6 | Bruker Biospin | www.bruker.com |
| AutoDock 4.2 | The Scripps Research Institute | http://autodock.scripps.edu/ |
| AutoDockTools 1.5.6 | The Scripps Research Institute | http://autodock.scripps.edu/resources/adt |
| Synergy Finder | Institute for Molecular Medicine<br>Finland | https://synergyfinder.fimm.fi/ |
| Prism | GraphPad | https://www.graphpad.com/ |

### Data Deposition

BioMagResDB accession number: 50568 Title: “^1^H and ^15^N assignments for 14-residues peptide that is cleaved by SARS-CoV-2 M^pro^.”

BioMagResDB accession number: 50569. Title: “^1^H and ^15^N assignments for 14-residue peptide after cleavage by SARS-CoV-2 M^pro^”

### Resource Availability

#### Lead Contact

Further information and requests for resources and reagents should be directed to and will be fulfilled by the designated contacts Gaetano T. Montelione.

#### Materials Availability

All materials generated in this study are available upon request.

### Method Details

#### Molecular Docking

We used the free open source *Autodock* suite (Morris et al., 2009). *AutoDockTools* (Sanner, 1999) was used for coordinate preparation, docking, and analysis of results. The computational docking program *AutoDock v4.2.6*. is based on an empirical free- energy force field and uses a search method based on Lamarckian genetic algorithm (Morris et al., 1998). Target protein coordinates were obtained from SARS-CoV-2 M^pro^ X-ray crystal structure (PDB id 6Y2G) (Zhang et al., 2020), and structural water was removed. Three-dimensional coordinates for ligand molecules were obtained from PDB (http://www.rcsb.org/) or from chemical structure databases, ChemSpider (http://www.chemspider.com/) and DrugBank (https://www.drugbank.ca/). Protein and ligand coordinates were then prepared using *AutoDockTools*; polar hydrogens were added to protein structures, and Gasteiger-Marsili empirical atomic partial charges were added to ligands. Torsional degrees of freedom (dihedral angles) were identified for each ligand. These data and parameters for each protein and ligand were saved as individual PDBQT files. In these studies, ligand dihedral angles were allowed to vary (except where stated otherwise), and all protein dihedral angles were kept rigid. The program *Autogrid* (Sanner, 1999) was used to prepare affinity maps for all atom types in the receptor and ligands. A grid of 56, 40 and 48 points in x, y and z direction, with a grid spacing of 0.375 Å was used to compute electrostatic maps. The grid center was placed on the center of the inhibitor 13b molecule in its complex with M^pro^ (PDB id 6Y2G) (Zhang et al., 2020). The Lamarckian genetic algorithm (LGA) method was used for sampling ligand binding conformation (Morris et al., 1998), with the following LGA parameters: 150 individuals in population; 2,500,000 energy evaluations; 27,000 maximum number of generations; and with mutation and crossover rates of 0.02 and 0.08, respectively. A maximum of 300 iterations per local search was used. The calculations were repeated for 100 docking simulations for each complex. For a comparative analysis, docking simulations of the α-ketoamide inhibitor 13b (Zhang et al., 2020) were also performed using the same protocol used for docking the protease inhibitor drugs. All docking simulations were analyzed using the *AutoDockTools.* Atomic coordinates for best scoring conformation obtained in each docking simulation, for each drug-protein complex, were saved in PDB format for analysis. These protein-ligand complexes were analyzed in detailed using open source *PyMol* molecular visualization tool (DeLano, 2009) and fully automated *Protein-Ligand Interaction Profiler* (Salentin et al., 2015) (https://projects.biotec.tu-dresden.de/plip-web/plip).

PL^pro^ inhibitor complexes in the PDB revealed that the BL2 loop present at the entrance of active site adopts significantly different conformations depending on the size of the inhibitor bound to the PL^pro^. This plasticity in the BL2 loop suggests an induced fit mechanism of ligand binding to PL^pro^ active site. To avoid a closed conformation of the BL2 loop found in protein ligand complex we chose SARS-CoV-2 PL^pro^ X-ray crystal structure in its apo form (PDB id 6W9C) as the target to dock HCV protease inhibitors. The docking protocol was same as above, except a larger grid of size 56, 56, and 58 points in the x, y and z direction respectively was used to compute electrostatic maps for PL^pro^ target. For a comparative analysis, docking simulations of PL^pro^ inhibitor GRL0617 (PDB id 7CJM) were also performed using the same protocol.

#### M^pro^ expression and purification

The full length SARS-CoV-2 M^pro^ gene was ordered from GenScript USA Inc. in pGEX- 6P-1 vector, as previously described (Zhang et al., 2020). This expression vector is designated GTM_COV2_NSP5_001. This plasmid, expressing SARS-CoV-2 M^pro^ as a self-cleaving (using its native cleavage site) GST-fusion were transformed into competent *E. coli* BL21(DE3) cells. A single colony was picked and inoculated in 2 mL LB supplemented with 0.1 mg/ml Ampicillin at 37 °C and 225 rpm. The 2 mL inoculum was added to 1L LB broth with 0.1 mg/mL Ampicilin. The cells were allowed to grow to an optical density of 0.6 at 600 nm at 37 °C and 225 rpm, and induced with 1 mM IPTG. The induced cells were incubated overnight at 18 °C and 225 rpm. The cells were harvested and resuspended in lysis buffer (20 mM Tris pH 8.0, 300 mM NaCl) and then lysed by sonication. The cell debris was removed by centrifugation at 14000 rpm for 40 mins. The supernatant was then added to a Ni-NTA column. The Ni-NTA column was then washed with wash buffer (20 mM Tris pH 8.0, 300 mM NaCl, 10 mM Imidazole) and eluted with 20 mM Tris pH 8.0, 100 mM NaCl, 100 mM imidazole. The elution fractions that contained M^pro^ were simultaneously buffer exchanged (20 mM Tris pH 8.0, 100 mM NaCl, 1 mM EDTA, 1 mM DTT) and concentrated using a swinging bucket centrifuge at 4000 rpm. The final M^pro^ concentration was 23.3 µM, determined by absorbance at 280 nm using a calculated extinction coefficient of 33,640 M^-1^ cm^-1^. Homogeneity was validated by SDS-PAGE (> 95% homogeneous) and the construct was validated by MALDI-TOF mass spectrometry.

#### M^pro^ proteolysis inhibition assays

The proteolysis of substrate KTSAVLQ/SGFRKME was studied using Fluorescence Resonance Energy Transfer (FRET) and Nuclear magnetic Resonance (NMR) assays. For the FRET assay the substrate was labeled with Dabcyl and Glu-Edans FRET pair on the N and C-termini of the peptide, respectively, as described elsewhere(Ma et al., 2020). Both, labeled and unlabeled substrates were ordered from GenScript USA Inc. For all assays, the specific activity of the enzyme was checked at the beginning and end of the data collection session, or back-to-back with each measurement, in order to avoid spurious results due to enzyme inactivation during a measurement session.

All M^pro^ proteolysis assays were carried out in Reaction Buffer containing 20 mM HEPES pH 6.5, 120 mM NaCl, 0.4 mM EDTA and 3 mM TCEP. Additionally, 1 mg/mL BSA was added to the buffer for FRET assays. For the FRET assay, 10 nM M^pro^ was incubated with 20 µM of HCV drugs. The reaction was initiated by addition of 20 µM FRET substrate and monitored for 2 hours on an Infinite M1000 TECAN plate reader, exciting at 360 nm and detecting donor emission at 460 nm. The initial velocity of the reaction was calculated as the slope obtained from linear fits of emission intensity vs time plots for the first 15 minutes of the reaction. All FRET data were analyzed and plotted for initial velocity on Microsoft Excel.

In the FRET assays, the percent proteolytic activity in presence of each drug was calculated as a ratio of initial velocity in presence if inhibitor (*v_i_*) to initial velocity in absence of inhibitor (*v_i0_*) i.e. *v_i_ / v_i0_*. A histogram plot of *v_i_ / v_i0_* for each inhibitor was used to compare relative inhibition activities. All FRET proteolysis reaction curves were measured twice, and the uncertainty in *v_i_ / v_i0_*, estimated from the standard deviation of 2 independent measurements, is shown as error bars.

For IC_50_ measurements of HCV inhibitors BOC, NAR and TEL, 10 nM of M^pro^ was incubated with a range of inhibitor concentrations in the same Reaction Buffer described above. The inhibitor concentration ranges were 0.1 – 200 μM for BOC, 0.1 to 100 μM for NAR, and 0.2 to 100 μM for TEL. The reaction was initiated by adding 20 µM FRET substrate and monitored for 1 hr. Each measurement was repeated three times. The % inhibition at each inhibitor concentration was calculated as:

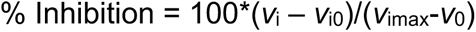

where v_i_ = initial velocity at a given inhibitor concentration

*v*_i0_ = initial velocity in absence of inhibitor

*v*_imax_ = initial velocity at maximum inhibition

The % inhibition was plotted as a function of inhibitor concentration to obtain a dose-response curve using Graphpad Prism 8 software. IC_50_ was calculated from fitting to the equation:

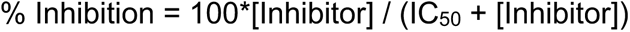

For the ^1^H NMR proteolysis assay the reaction was performed at 100 nM M^pro^ in the same assay buffer described above, along with 5% D_2_O and 50 µM HCV inhibitors dissolved in d_6-_DMSO (for the control experiments where no inhibitor is added, the same quantity of d_6-_DMSO was added). 50 µM of unlabeled substrate was added and immediately transferred to a 5-mm NMR tube. The final volume of each reaction mixture was 600 µL. The NMR tube was quickly placed in a 600 MHz Bruker Avance II spectrometer equipped with a 5-mm TCI cryoprobe, equilibrated at 298K. The homogeneity of the magnetic field was adjusted by gradient shimming on the z-axis and in each case an array of 24 ^1^H experiments was acquired with 1D ^1^H NMR using excitation sculpting for water suppression. The probe had previously been tuned and matched with a sample of similar composition. The delay between initiation of the reaction and starting acquisition was ∼ 5 mins for most of the reaction conditions. The duration of each NMR experiment was also taken into account to obtain accurate time values. All ^1^H spectra were acquired, processed, and analyzed in Bruker TopSpin 3.6.2 software. The regions of interest were integrated, and the values obtained were transferred for further analysis and plotting.

These ^1^H NMR spectra were used to monitor the evolution of substrate and product as a function of time. Resonance assignments (discussed below) of the cleaved and uncleaved KTSAVLQ/SGFRKME peptide identified the amide H^N^ resonances that were monitored during the reaction. The H^N^ resonances for amino-acid residues Phe-10 (uncleaved) and Gln-7 (cleaved) were used to quantify substrate utilization and product formation, respectively, during the reaction. The H^N^ preak intensity of residue Glu-14, which did not shift upon cleavage, was monitored as an internal control. The percent substrate cleavage in the presence of inhibitors at 30 min was calculated as a ratio of the H^N^ resonance integrals of Gln-7 in presence of inhibitor to the corresponding resonance integral with no inhibitor.

#### Amide ^1^H and ^15^N chemical shift assignments for M^pro^ peptide substrate

Chemical shift assignments of backbone amide ^1^H and ^15^N resonances in the 14- residue peptide KTSAVLQSGFRKME in 20 mM HEPES pH 6.5, 100 mM NaCl, 0.4 mM EDTA, 3 mM TCEP and 5% ^2^H_2_O were determined at 298 K using 2D COSY, TOCSY, and ^1^H-^15^N HSQC, along with 1D ^1^H NMR experiments, and referenced to internal DSS. Backbone amide ^1^H and ^15^N resonances were assigned for 12/14 and 10/14 residues in the uncleaved and cleaved peptide respectively, and deposited in the BioMagResDB as BMRB entries 50568 and 50569, respectively

#### PL^pro^ proteolysis inhibition assay

SARS-CoV-2 PL^pro^ enzyme was provided as a generous gift by Prof. John Hunt (Columbia University). The construct was produced in expression vector pET21_NESG with a C-terminal purification tag LEHHHHHH. Homogeneity (>95%) was validated by SDS-PAGE. The fluorogenic substrate zRLRGG/AMC was ordered from Bachem. All PL^pro^ proteolysis assays were carried out in buffer containing 50 mM HEPES, pH 7.5, 5 mM DTT, 1 mg/ml BSA. For the fluorescence assay, 20 nM of PL^pro^ was incubated with 20 µM of HCV drugs. 20 µM substrate was added and the reaction was monitored for 2 hrs using the Infinite M1000 TECAN plate reader compatible with Magellan ^TM^ software with filters for excitation at 360 nm and emission at 460 nm. The data points for the first 10 mins of the proteolysis reaction progression curves were used to calculate the initial velocity (*v_i_*) in presence and absence of inhibitors. The percent proteolytic activity in presence of each drug is calculate as a ratio of initial velocity in presence of inhibitor (*v_i_*) to initial velocity in absence of inhibitor (*v_i0_*), i.e. *v_i_ / v_i0_*. A histogram plot of *v_i_ / v_i0_* for each inhibitor was used to compare relative inhibition activities. All proteolysis reaction curves were measured twice, and the range in *v_i_ / v_i0_* is shown as error bars.

#### Cells and Viruses

Vero E6 (ATCC, CRL-1586) and HEK293T (ATCC, CRL-3216; kind gift of Dr. Viviana Simon), were maintained in DMEM (Corning) supplemented with 10% FB (Peak Serum) and Penicillin/Streptomycin (Corning) at 37°C and 5% CO_2_. hACE2-293T cells were generated for this study. Briefly, HEK293T cells were transduced with a lentiviral vector expressing human ACE2. Puromycin resistant cells with hACE2 surface expression were sorted after staining with AlexaFluor 647-conjugated goat anti-hACE2 antibodies. Cells were then single-cell-cloned and screened for their ability to support SARS-CoV-2 replication. All cell lines used in this study were regularly screened for mycoplasma contamination using the Universal Detection Kit (ATCC, 30-1012K). Cells were infected with SARS-CoV-2, isolate USA-WA1/2020 (BEI Resources NR-52281) under biosafety level 3 (BSL3) containment in accordance to the biosafety protocols developed by the Icahn School of Medicine at Mount Sinai. Viral stocks were grown in Vero E6 cells as previously described (Amanat et al., 2020) and were validated by genome sequencing.

#### Viral growth and cytotoxicity assays in the presence of inhibitors

2,000 Vero E6 or hACE2-293T cells were seeded into 96-well plates in DMEM (10% FBS) and incubated for 24 h at 37C, 5% CO2. Two hours before infection, the medium was replaced with 100 µl of DMEM (2% FBS) containing the compound of interest at concentrations 50% greater than those indicated, including a DMSO control. Plates were then transferred into the BSL3 facility and 100 PFU (MOI = 0.025) was added in 50 µl of DMEM (2% FBS), bringing the final compound concentration to those indicated. Plates were then incubated for 48 h at 37C. After infection, supernatants were removed and cells were fixed with 4% formaldehyde for 24 hours prior to being removed from the BSL3 facility. The cells were then immunostained for the viral NP protein (an in-house mAb 1C7, provided by Dr. Thomas Moran (MSSM) with a DAPI counterstain. Infected cells (488 nM) and total cells (DAPI) were quantified using the Celigo (Nexcelcom) imaging cytometer. Infectivity was measured by the accumulation of viral NP protein in the nucleus of the Vero E6 cells (fluorescence accumulation). Percent infection was quantified as [(Infected cells/Total cells) – Background] *100 and the DMSO control was then set to 100% infection for analysis. The IC_50_ and IC_90_ for each experiment were determined using the Prism (GraphPad Software) software. Cytotoxicity was also performed using the MTT assay (Roche), according to the manufacturer’s instructions. Cytotoxicity was performed in uninfected VeroE6 cells with same compound dilutions and concurrent with viral replication assay. All assays were performed in biologically independent triplicates. Remdesivir was purchased from Medkoo Bioscience inc. Time of addition experiments were performed using the same immunofluorescence-based assay with the following alterations: Vero E6 cells we infected with 8000 PFU (MOI of 2) of SARS-CoV-2, compound was added at different times relative to infection as indicated, and the infection was ended by fixation with 4% formaldehyde after 8 hours (single cycle assay).

#### Antiviral combination assay

Like previous antiviral assay, 2,000 Vero E6 cells were seeded into 96-well plates in DMEM (10% FBS) and incubated for 24 h at 37°C, 5% CO_2_. Two hours before infection, the medium was replaced with 100 µl of DMEM (2% FBS) containing the combination of HCV protease inhibitors and remdesivir following a dilution combination matrix. A 6 by 6 matrix of drug combinations was prepared in triplicate by making serial two-fold dilutions of the drugs on each axis, including a DMSO control column and row. The resulting matrix had no drug in the right upper well, a single drug in rising 2-fold concentrations in the vertical and horizontal axes starting from that well, and the remaining wells with rising concentrations of drug mixtures reaching maximum concentrations of the drugs at the lower left well. Plates were then transferred into the BSL3 facility and SARS-CoV-2 (MOI 0.025) was added in 50 µl of DMEM (2% FBS), bringing the final compound concentration to those indicated in the figures. Plates were then incubated for 48 h at 37 °C. After infection, cells were fixed with final concentration of 5% formaldehyde for 24 hours prior to being removed from the BSL3 facility. The cells were then immunostained for the viral NP protein (an in-house mAb 1C7, provided by Dr. Thomas Moran, MSSM) with a DAPI counterstain. Infected cells (AlexaFluor 488) and total cells (DAPI) were quantified using the Celigo (Nexcelcom) imaging cytometer. Infectivity was measured by the accumulation of viral NP protein in the nucleus of the Vero E6 cells (fluorescence accumulation). Percent infection was quantified as [(Infected cells/Total cells) – Background] *100 and the DMSO control was then set to 100% infection for analysis. The combination antiviral assay was performed in biologically independent triplicates. The apparent IC_90_ for each combination in the matrix was determined using the Prism (GraphPad Software) software. The IC_90_ for HCV drugs and remdesivir were calculated for each drug treatment alone and in combination. This combination data was analyzed using SynergyFinder by the ZIP method (Ianevski et al., 2020), and combination indices were calculated as previously described (Amanat et al., 2020).

## Supplementary Information

### Supplementary Tables

**Supplementary Table S1.** SARS-CoV-2 M^pro^ and HCV 3C/4A Protease Inhibitors

| Inhibitor<br>(Trade Name) | Identifier<br>of<br>Protease<br>Inhibitor | Database ID<br>of Protease<br>Inhibitor<br>Structure | AutoDock Score<br>(kcal/mol)<br>Lowest<br>“Energy” |  |
| --- | --- | --- | --- | --- |
|  |  |  | M<sup>pro</sup> | PL<sup>pro</sup> |
| **<u>SARS-CoV-2 M<sup>pro</sup> Inhibitor</u>** |  |  |  |  |
| α-ketoamide inhibitor 13b<br>lowest “energy” pose:<br>pose most similar to X-ray structure: | 06K | 6Y2G<sup>a</sup> | -9.17<br>-9.03 | Not<br>Applicable |
| **<u>SARS-CoV-2 PL<sup>pro</sup> Inhibitor</u>** |  |  |  |  |
| GRL0167<br>lowest “energy” pose |  | 7CJM<sup>a</sup> | Not<br>Applicable | -7.54 |
| **<u>HCV NSP3/4A</u>**<br>**<u>Protease Inhibitor Drugs</u>** |  |  |  |  |
| Vaniprevir | VAN | 3SU3<sup>c</sup> | -10.95 | -7.85 |
| Simeprevir | SIM | 3KEE<sup>c</sup> | -10.75 | -8.12 |
| Paritaprevir | PAR | 32700634<sup>b</sup> | -10.71 | -10.30 |
| Danoprevir | DAN | 3M5L<sup>c</sup> | -9.99 | -8.48 |
| Narlaprevir | NAR | 3LON<sup>c</sup> | -9.80 | -5.56 |
| Grazoprevir | GRZ | 3SUD<sup>c</sup> | -9.71 | -8.10 |
| Glecaprevir | GLE | 35013015<sup>b</sup> | -9.51 | -8.30 |
| Boceprevir | BOC | 10324367<sup>e</sup> | -9.13 | -5.56 |
| Telaprevir | TEL | 3SV6<sup>c</sup> | -9.05 | -6.57 |
| Asunaprevir | ASU | 4WF8<sup>c</sup> | -8.37 | -6.57 |

(a) Atomic coordinates for the inhibitor taken from listed PDB id. (b) Atomic coordinates for the inhibitor were taken from the ChemSpider database. (c) Atomic coordinates for the inhibitor were taken from the PDB coordinates of the corresponding complex of the inhibitor bound to HCV 3C/4A protease. (d) Mavyret (or Maviret) is a multidrug formulation including glecaprevir and pibrentasvir. (e) Atomic coordinates for the inhibitor were taken from the Pubchem database.

**Supplementary Table S2.** Key Interactions within the SARS-CoV-2 M^pro^ Substrate Binding Cleft

| Inhibitor | Identifier of<br>Protease<br>Inhibitor | Atoms Forming<br>Key Hydrogen Bonds | Residues Forming Key<br>Hydrophobic<br>Interactions |
| --- | --- | --- | --- |
| **<u>SARS-CoV-2 M<sup>pro</sup> Inhibitor</u>** |  |  |  |
| α-ketoamide<br>inhibitor 13b<br>(X-ray crystal structure<br>PDB id 6Y2G) | 06K | His41(N<sup>ε</sup>), Gly143(N), Ser144(N),<br>Cys145(N), His163(N<sup>ε</sup>), His164(O),<br>Glu166(O<sup>ε</sup>) | Asn142, Met165, Asp187,<br>Gln189 |
| α-ketoamide<br>inhibitor 13b<br>(no dihedral rotations) | 06K | His41(N<sup>ε</sup>), Gly143(N), Ser144(N, O<sup>γ</sup>),<br>Cys145(N), His163(N<sup>ε</sup>), Glu166(N, O<sup>ε</sup>) | Asn142, Met165, Pro168,<br>Gln189 |
| α-ketoamide<br>inhibitor 13b<br>(lowest “energy” pose) | 06K | Thr26(N), His41(N<sup>ε</sup>),<br>Gly143(N), Cys145(N) | His41, His164,<br>Met165, Glu166, Phe181 |
| α-ketoamide<br>inhibitor 13b<br>(pose most similar to X-ray<br>structure) | 06K | His41(N<sup>ε</sup>), Gly143(N),<br>Glu166(N, O) | Thr25, Phe140, Met165,<br>Leu167, Pro168, Gln189,<br>Gln192 |
| **<u>HCV NSP3/4A</u>**<br>**<u>Protease Inhibitor Drugs</u>** |  |  |  |
| Vaniprevir | VAN | His41(N<sup>ε</sup>), Asn142(O<sup>δ</sup>), Gly143(N),<br>Glu166(N) | Thr25, Phe140, Asn142,<br>Glu166, Pro168 |
| Simeprevir | SIM | Thr25(O<sup>γ</sup>), His41(O), Cys44(N),<br>Glu166(C<sup>β</sup>) | Thr24, Met165, Glu166,<br>Gln189 |
| Paritaprevir | PAR | Asn142(O<sup>δ</sup>), Gly143(N), Ser144(N),<br>Cys145(N), His163 (N<sup>δ</sup>), Glu166(O,N) | Leu27, Gln189, Gln192 |
| Narlaprevir | NAR | His41(N<sup>ε</sup>), Asn142(N, O<sup>δ</sup>), Gly143(N),<br>Ser144(N), Cys145(N), Glu166 (N,O) | Thr25, Leu27, Met165,<br>Leu167, Gln192 |
| Danoprevir | DAN | Thr26(N), Asn142(O<sup>δ</sup>), Gly143(N),<br>His163(N<sup>ε</sup>), Glu166(N) | Thr25, Leu27, Asn142,<br>Glu166 |
| Grazoprevir | GRZ | Gly143(N), Ser144(N), Cys145(N),<br>Glu166(N) | Thr25, Leu27, Met165,<br>Pro168, Gln189 |
| Glecaprevir | GLE | His41(N<sup>ε</sup>), Asn142(N), Gly143(N),<br>Glu166(N) | Thr25, Leu141, Met165,<br>Glu166 |
| Boceprevir | BOC | His41(N<sup>ε</sup>), Gly143(N), Ser144(N),<br>Glu166(N, O) | His41, Phe140, Met165,<br>Glu166 Leu167 Pro168,<br>Asp187 |
| Telaprevir | TEL | His41(N<sup>ε</sup>), Gly143(N), Ser144(N),<br>Cys145(N), Glu166(N) | Thyr25, Gln189 |
| Asunaprevir | ASU | His41(N<sup>ε</sup>), Asn142(N), Glu166(N),<br>Arg188(O) | Thr25, Met165, Glu166,<br>Gln189 |

## Supplementary Figures

**Fig. S1:**
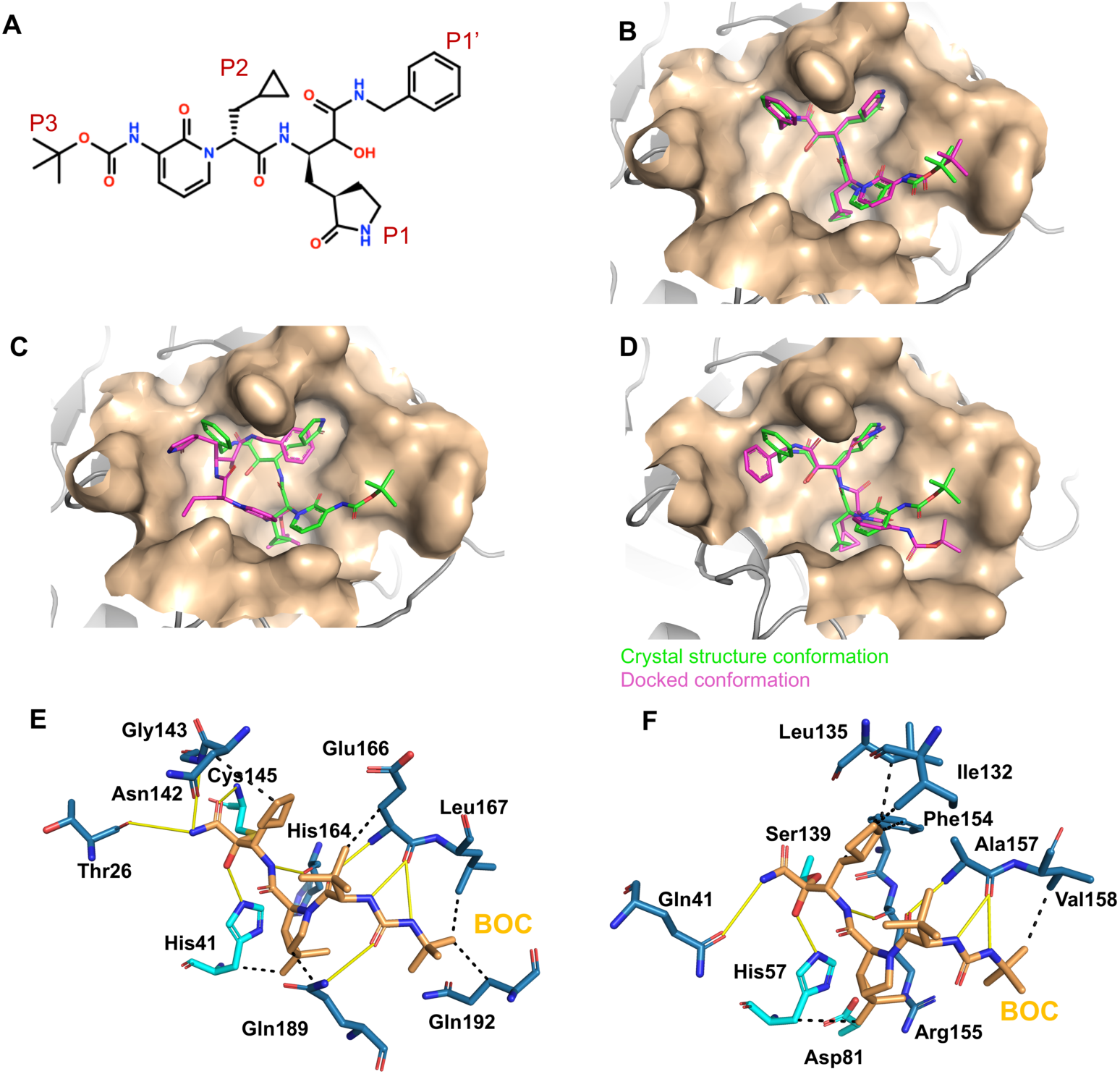
Interaction of a-ketoamide inhibitors 13b and boceprevir to SARS-CoV-2 M^pro^. (A) Chemical structure of M^pro^ inhibitor α-ketoamide inhibitor 13b. (B) Lowest “energy” *AutoDock* pose using a rigid conformation of α-ketoamide inhibitor 13b, to match the ligand conformation in X-ray crystal structure of the complex (score = -7.19 kcal/mol). (C) Lowest “energy” *AutoDock* pose observed among 100 docking simulations (score = -9.17 kcal/mol). (D) The low “energy” *AutoDock* pose (*AutoDock score* = -9.03 kcal/mol) of 13b that is most similar to the conformation seen in the crystal structure. In panels B-D, M^pro^ is shown in surface representation, X-ray crystal structure of α-ketoamide inhibitor 13b bound in the active site of M^pro^ (green sticks, PDB id 6Y2G), and the predicted *AutoDock* conformation (magenta sticks). (E) Details from the X-ray crystal structure of the SARS-CoV-2 M^pro^ protease - BOC complex (PDB id 6WNP). (F) Details from the X-ray crystal structure of the HCV NS3/4A protease - BOC complex (PDB id 2OC8). Boceprevir forms a covalent bond with the catalytic Ser or Cys residues, respectively, inhibiting both of these proteases. In both of these boceprevir – protease complex structures, remarkable similarity is observed between the predicted and observed binding poses and protein-inhibitor hydrogen bond networks. In the complex with HCV protease, boceprevir forms hydrogen bonds with side chains of residues Gln41 and His57 and with backbone atoms of Gly137, Ser139, Arg155 and Ala157. The corresponding residues equivalents (based on structural superimposition) of these residues in M^pro^, Thr26, His41, Gly143, Cys145, His164 and Glu166, also form hydrogen bonds with boceprevir. The sidechain of residue Gln189 of SARS-CoV-2 M^pro^ forms an additional hydrogen-bond with boceprevir.

**Fig. S2:**
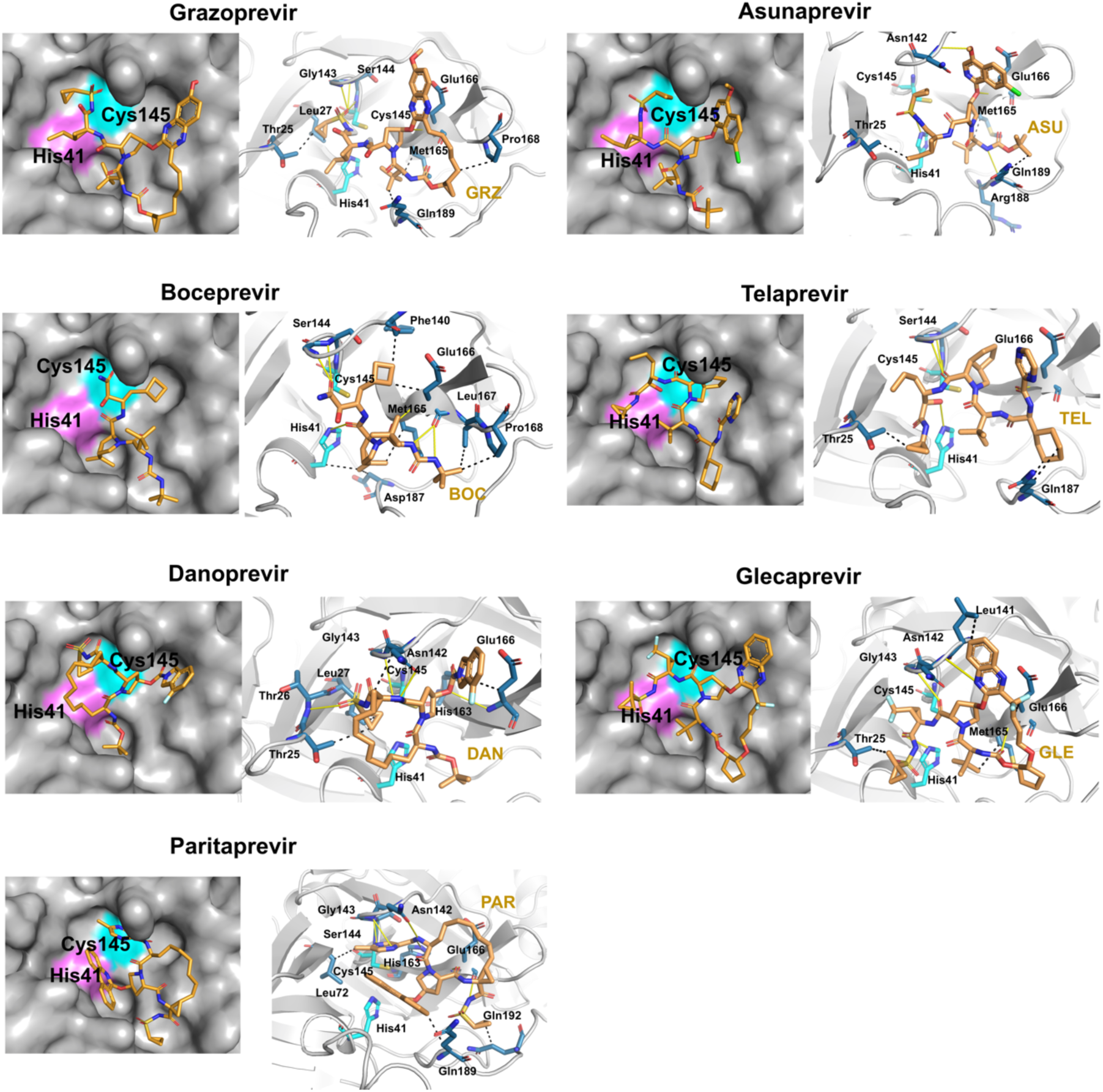
Docking of HCV protease NS3/4A inhibitor drugs to SARS-CoV-2 M^pro^. Right panels - Lowest “energy” *AutoDock* pose of these HCV protease inhibitors (orange sticks) into the SARS-CoV-2 M^pro^ active site. M^pro^ is shown in surface representation (gray), surfaces of catalytic residues His41 (magenta) and Cys145 (cyan) are highlighted. Left panels – Details of atomic interactions in the lowest “energy” *AutoDock* poses of these HCV protease inhibitors with M^pro^. Hydrogen bonds and hydrophobic interactions between the drug and the enzyme are shown with yellow solid lines and black dashed lines, respectively. Sidechains of catalytic residues (cyan) His41 and Cys145 are shown as sticks and labeled, along with other protein residues (marine blue) that form key interactions with these drugs.

**Fig. S3:**
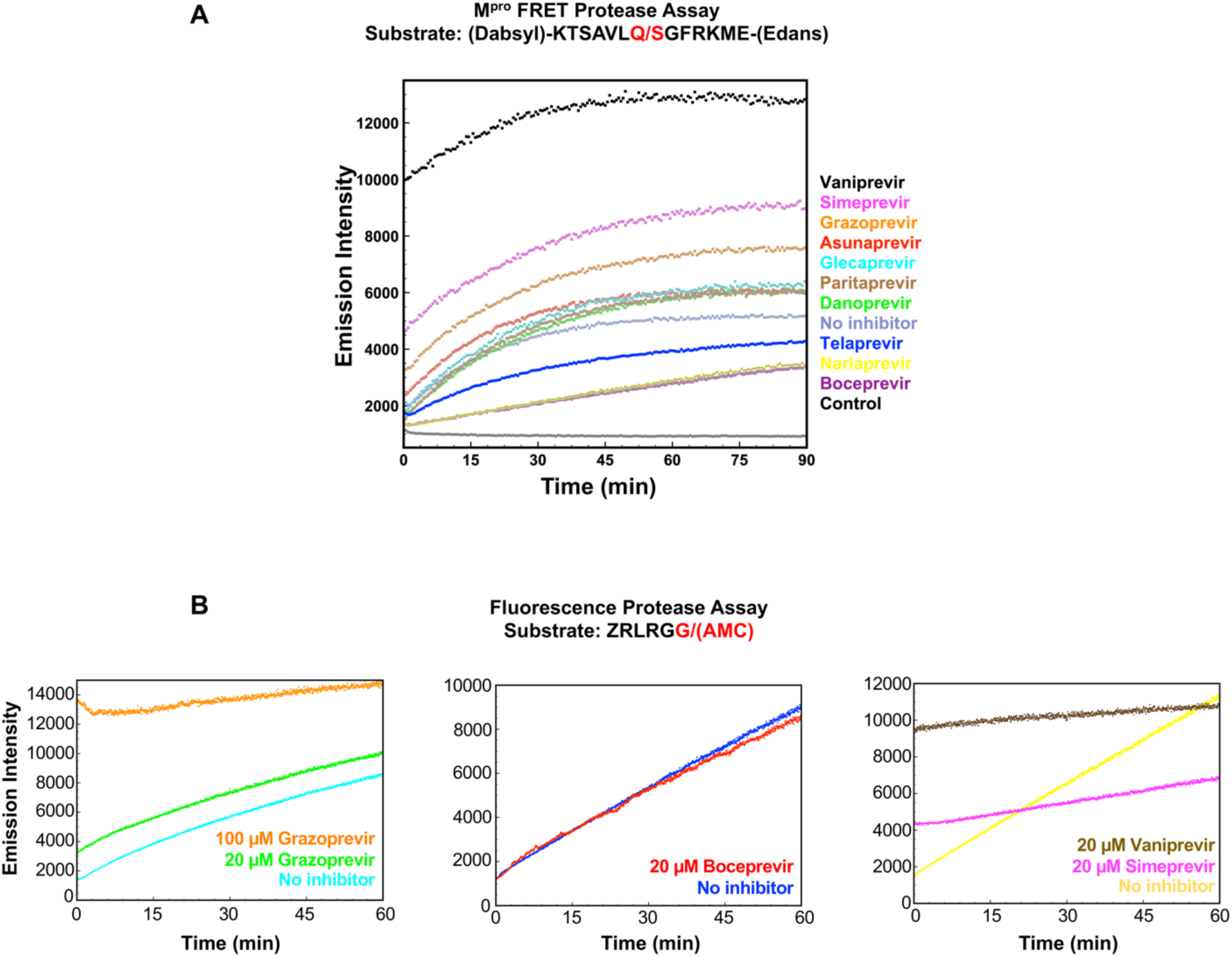
Proteolytic cleavage progression curves. (A) Progression kinetics for M^pro^ were monitored in a FRET assay where substrate was labelled with Dabcyl and Edans FRET pair on N and C termini respectively. (B) Progression kinetics for PL^pro^ were monitored in a florescence assay where substrate was labelled with a fluorogenic dye AMC. For both proteases, reaction progressions were monitored in presence and absence of HCV NS3/4A protease inhibitors for 60-90 mins, exciting at 360 nm and detecting donor emission at 460 nm.

**Fig. S4.**
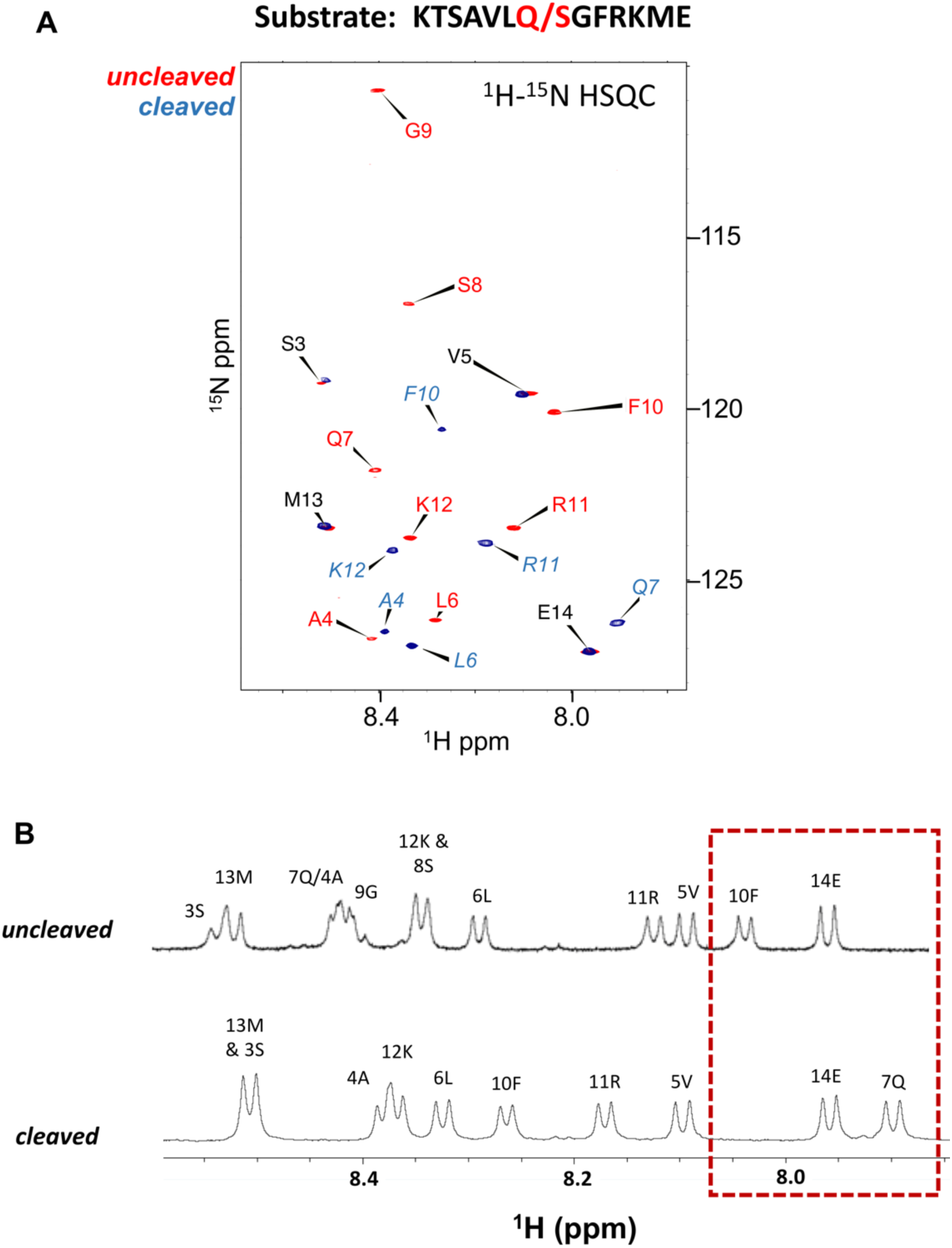
Chemical shift assignments of backbone amide protons in uncleaved and cleaved M^pro^ peptide substrate. (A) Overlay of 2D ^1^H-^15^N HSQC spectra for uncleaved (red) and cleaved (blue) peptides (B) 1D ^1^H spectra for cleaved and uncleaved peptides. Both spectra show changes in chemical shifts for some amino acids indicating proteolytic cleavage. These chemical shifts were determined at 25 °C using 2D COSY, TOCSY, and ^1^H-^15^N HSQC, along with 1D ^1^H NMR experiments.

**Fig. S5.**
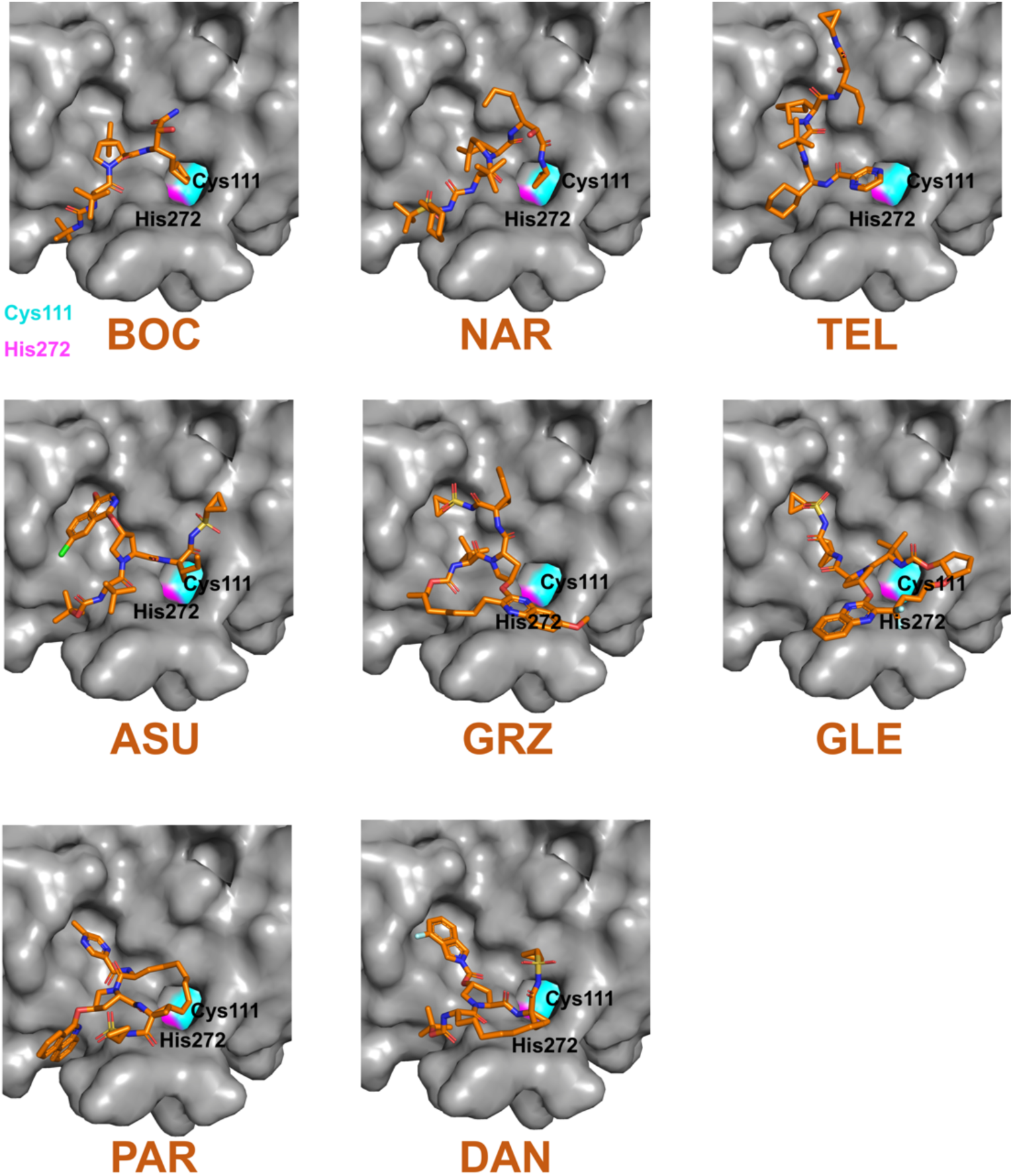
Docking of HCV protease NS3/4A inhibitor drugs to SARS-CoV2 PL^pro^. Lowest “energy” *AutoDock* pose of HCV protease NS3/4A inhibitors (orange sticks) into the SARS-CoV- 2 PL^pro^ active site are shown. PL^pro^ is shown in surface representation (gray), surfaces of catalytic residues His41 (magenta) and Cys145 (cyan) are highlighted.

**Fig. S6:**
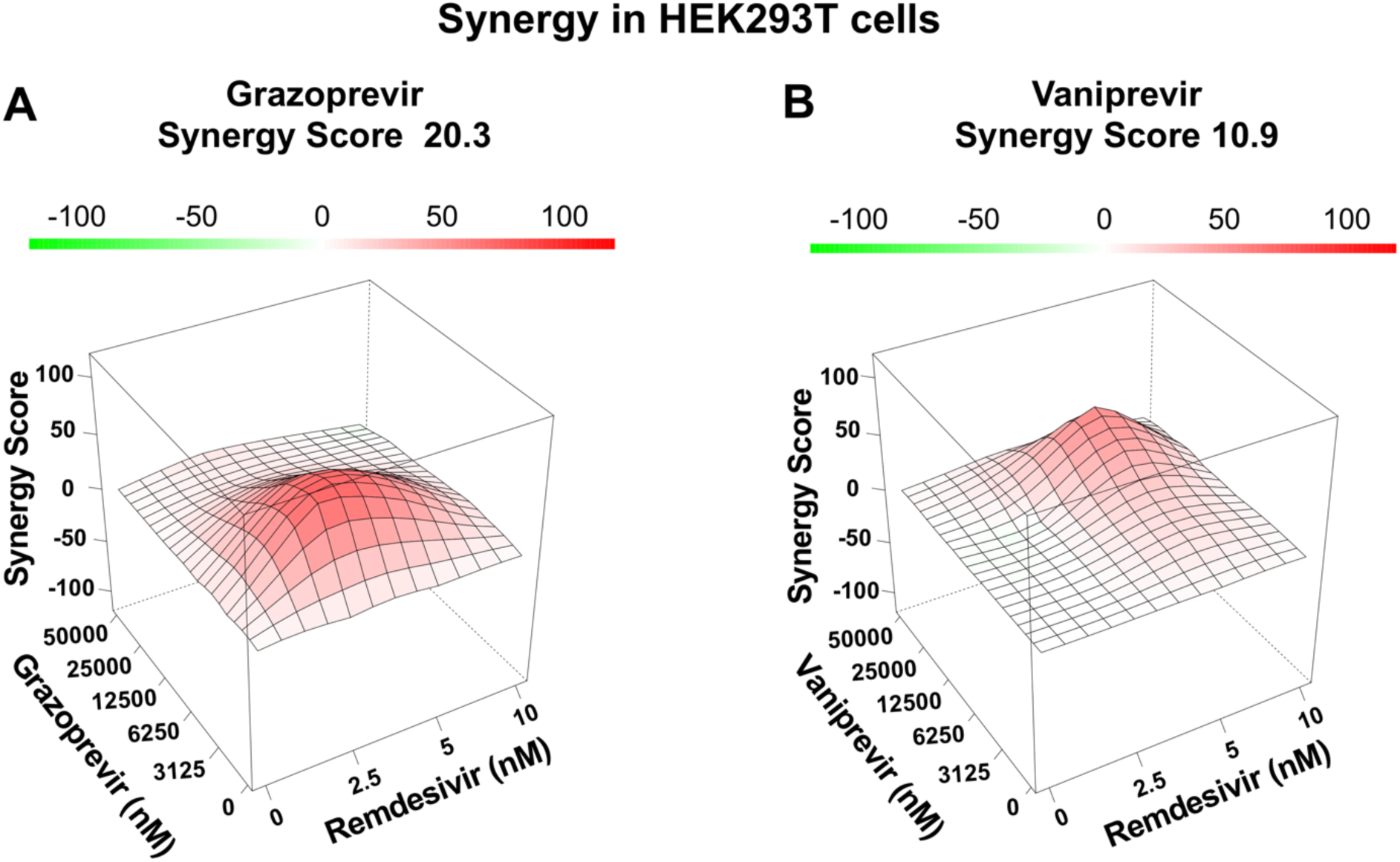
HCV inhibitors are synergistic with remdesivir in antiviral combination assay. Human HEK293T cells were infected with SARS-CoV-2 in presences of two compounds titrated against each other in 2-fold serial dilutions and viral replication was determined using our immunofluorescence-based assay. Both (A) grazoprevir and (B) vaniprevir have positive ZIP synergy score (Ianevski et al., 2020), indicative of synergy between these drugs and remdesivir in the combination assays. However, the synergy score for grazoprevier (+20.3) clearly indicates moderate synergy (Ianevski et al., 2020), while the score for vaniprevir (+10.9) indicates very weak synergy or an additive effect (Ianevski et al., 2020).

**Fig. S7:**
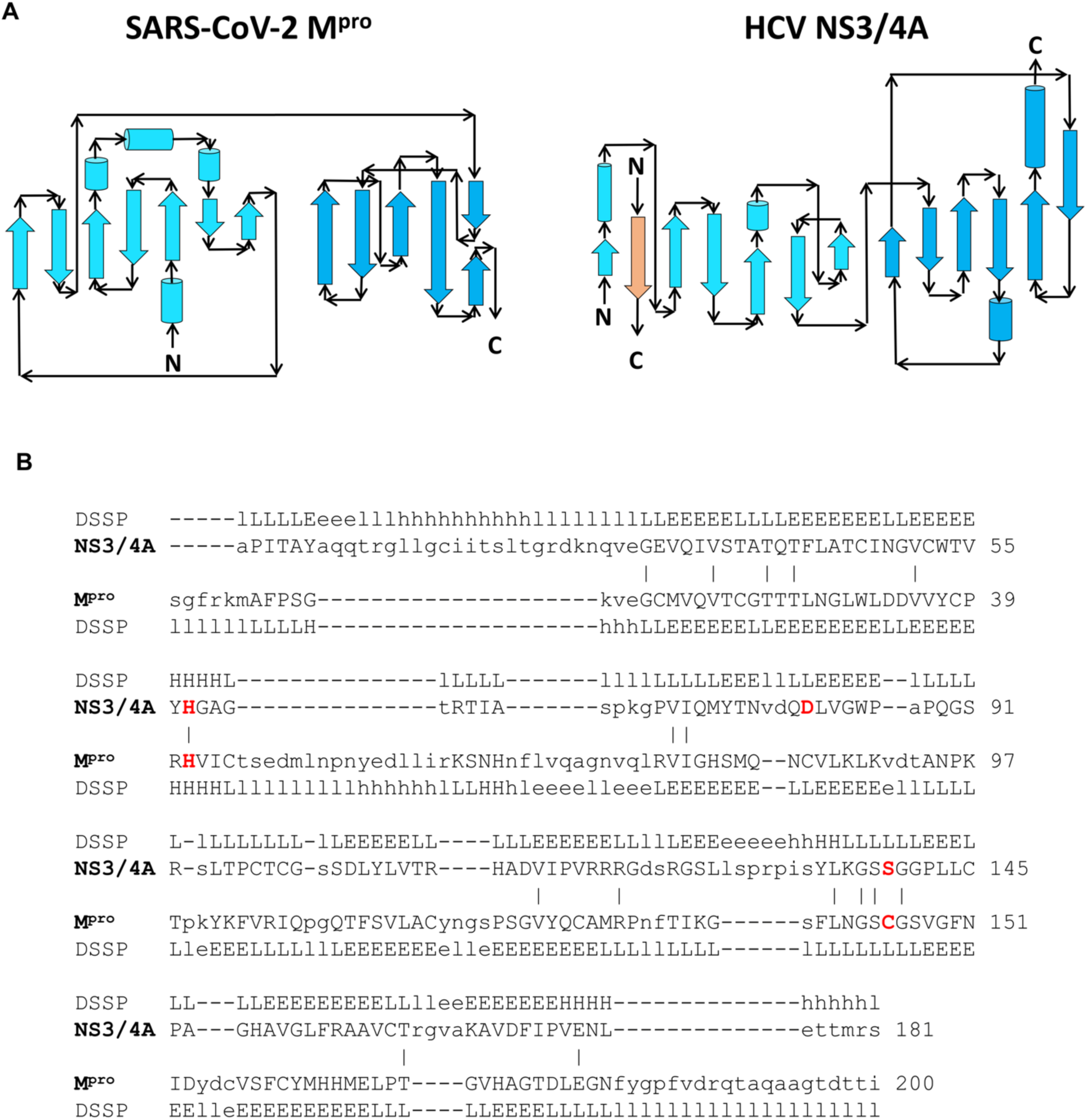
Comparison of fold topology diagrams and structure-based sequence alignments of SARS-CoV-2 M^pro^ and HCV NS3/4A proteases. (A) Fold topology diagrams demonstrate similar fold architectures but different fold topologies. (B) The DALI server provides a structure-based sequence alignment of HCV NS3/4A (HCV) and SARS-CoV-2 M^pro^ (M^pro^). Catalytic residues of HCV NS3/4A (His57, Asp81 and Ser139) and SARS-CoV-2 M^pro^ (His41 and Cys145) are highlighted in bold red. The structure-based alignment results in alignment of key catalytic residues His41 and Cys145 of the SARS-CoV-2 M^pro^ with His57 and Ser139 of HCV NS3/4A protease, and some other substrate binding cleft residues. Three-state secondary structure definitions by DSSP (H=helix, E=sheet, L=coil) are shown for each amino acid sequence. Structurally equivalent residues are in uppercase, structurally non-equivalent residues (e.g. in loops) are in lowercase. Identical amino acid are marked by vertical bars.

## Supplementary Results

### Validation of *AutoDock* protocols with inhibitor 13b

In the 1.75-Å X-ray crystal structure of the 13b-M^pro^ complex (Zhang et al., 2020), 13b makes hydrogen-bonded interactions with backbone amides of key residues, Gly143, Ser144 and Cys145, in the canonical oxyanion hole of the active site. To estimate the docking score for the pose observed in the X-ray crystal structure (Zhang et al., 2020), we first carried out docking using a modified protocol in which the dihedral angles of 13b were fixed to the values observed in the crystal structure. The lowest energy pose obtained with this protocol matches the crystal structure almost exactly (Supplementary Figure S1B), with *AutoDock* binding energy of -7.19 kcal/mol. Next, we assessed the docking protocol using flexible ligand dihedral angles, as was used for all the other HCV protease inhibitors. The best-scoring docked conformation (Supplementary Figure S1C), has an *AutoDock* binding energy of -9.17 kcal/mol. Because the experimentally-determined pose is often not the one with the lowest docking energy, but rather is found among other highly-ranked poses (Kolb et al., 2009; Kolb and Irwin, 2009), we also examined other low energy poses. One such pose, which has a slightly higher binding energy of -9.03 kcal/mol, is almost identical to the pose observed in the crystal structure (Supplementary Figure S1D). Hence, binding poses very similar to that observed in the crystal structure are indeed included among the low energy poses generated by our *AutoDock* protocol.

### Detailed descriptions of *AutoDock* lowest-energy binding poses observed for ten HCV protease inhibitors bound to M^pro^

**Asunaprevir** (ligand id ASU)

Molecular docking studies show that asunaprevir binds to the substrate binding cleft of M^pro^ with a predicted binding energy of -8.37 Kcal/mol (Figure 1D and Table S1). The key M^pro^ atoms involved in hydrogen bonding with ASU are the side chain N^ε^ atom of catalytic His41, backbone N atom of residues Asn142, Glu166 and Arg188 (Figure S2). In addition, residues Thr25, Met165, Glu166 and Gln189 contribute towards binding energy through hydrophobic interactions. In this binding conformation ASU does not interact with the residues forming the oxyanion hole and catalytic residues Cys145 that resides deep in the binding pocket. ASU rather binds in a conformation that can occlude the access of catalytic Cys145 by the polyprotein substrate of the protease.

**Boceprevir** (ligand id BOC)

Boceprevir is an alpha-ketoamide inhibitor like 13b and is therefore capable of covalently inhibiting the protease. In our docking studies BOC binds in the substrate binding cleft of M^pro^ with a predicted binding energy of -9.13 kcal/mol (Figure 1D and Table S1). The lowest scoring docking pose of BOC forms hydrogen bonds with side chain N^ε^ of catalytic His41 and backbone atoms of Gly143, Ser144 and Glu166 (Figure S2). Hydrophobic interactions with residues His41, Phe140, Met165, Glu166, Leu167, Pro186 and Asp187 further stabilized the ligand in the binding cavity. The α-ketoamide group is oriented towards residue Cys145 of the protease, with a distance between the S atom of Cys145 and C of BOC of 3.8 Å.

**Danoprevir** (ligand id DAN)

The lowest scoring pose for danoprevir in the substrate binding cleft of M^pro^ has a predicted binding energy of -9.99 kcal/mol (Figure 1D and Table S1). The catalytic residues His41 and Cys145 do not directly interact with DAN with either hydrogen bonds or hydrophobic interactions. Apparently because of its large size and macrocyclic nature, DAN cannot access these deeper residing catalytic residues. DAN occupies the oxyanion hole of the protease, forming hydrogen bonds with Asn142 sidechain O^δ^ and Gly143 backbone amide N atom (Figure S2). In addition, backbone amide N atoms of Thr26 and Glu 166, and sidechain N^ε^ of His163 also form hydrogen bonds with DAN. The pose is also stabilized by hydrophobic interactions involving residues Thr25, Leu27, Asn142 and Glu166.

**Glecaprevir** (ligand id GLE)

The predict binding energy of glecaprevir docked to the substrate binding cleft of M^pro^ is -9.51 kcal/mol (Figure 1D and Table S1). The catalytic residue Cys145, which is deeper in the binding pocket of M^pro^, is not accessible to GLE as observed for other macrocyclic HCV inhibitors (Figure S2). GLE forms hydrogen bonds with the sidechain N^ε^ atom of catalytic His41 along with backbone N atom of residues Asn142, Gly143 and Glu166. Residues that form hydrophobically contribute towards GLE binding are Thr25, Leu141, Met165, Glu166.

**Grazoprevir** (ligand id GRZ)

The lowest scoring pose of grazoprevir in the substrate binding cleft of M^pro^ has a predicted binding energy of -9.71 kcal/mol (Figure 1D and Table S1). GRZ is a macrocyclic compound and therefore occupies more extensive surface of the binding pocket when compared to *α*-ketoamide inhibitor 13b and boceprevir. The catalytic residue His41 does not directly interact with GRZ with either hydrogen bonds or hydrophobic interactions. GRZ forms hydrogen bonds with backbone amide N atoms of Gly143, Ser144, catalytic Cys145 occupying the oxyanion hole of the protease. It is also observed to forms hydrogen bond with the residues Glu166 (Figure S2). This binding mode of GRZ also includes extensive hydrophobic interactions with the protease, involving residues Thr25, Leu27, Met165, Pro168 and Gln189 that contribute to the stabilization of the GRZ-M^pro^ complex.

**Narlaprevir** (ligand id NAR)

The best scoring pose of narlaprevir has a predicted binding energy of -9.80 Kcal/mol (Figure 1D and Table S1). The NAR-protease interaction (Figure S2) is stabilized by a number of hydrogen bonds formed with N^ε^ atom of His41, side chain O and backbone N atoms of Asn142, backbone N atoms of Gly143, Ser144, Glu166, and catalytic Cys145. Backbone carbonyl atom of Glu166 also forms two hydrogen bonds with N atoms of NAR. Residues contributing to hydrophobic interactions are Thr25, Leu27, Met165, Leu167, Gln192. Although the α-ketoamide group of NAR is oriented towards residue Cys145 of the protease, the distance between the S atom of Cys145 and C of NAR is 4.4 Å, which is larger than the distance in the 13b-M^pro^ crystal structure.

**Paritaprevir** (ligand id PAR)

Paritaprevir is a macrocyclic compound, and the predicted binding energy of the lowest scoring pose for PAR is -10.71 kcal/mol (Figure 1D and Table S1). In this pose PAR occupies the oxyanion hole of the protease and forms hydrogen bonds with backbone N atoms of GLy143, Ser 144, Cys145 (Figure S2). PAR can for hydrogen bonds with sidechain O^δ^ atom of Asn142, N^δ^ of His 163 and carbonyl O atom of Glu166 as well. This pose also appears to be stabilized by hydrophobic interactions involving residues Leu27, Gln189 and Gln192.

**Simeprevir** (ligand id SIM)

The predicted binding energy of the best scoring complex of simeprevir is -10.75 kcal/mol (Figure 1D and Table S1). In this pose, SIM sits in the binding cavity differently than inhibitor 13b (Figure S2). The key atoms involved in hydrogen bonding with SIM are Thr25 sidechain hydroxyl OH, His45 backbone carbonyl O, Cys44 backbone N, and Glu166 backbone N. In addition, sidechains of residues Thr24, Met165, Glu166, and Gln189 form hydrophobic interactions with SIM. SIM does not occupy the binding site near the catalytic residue Cys145; rather, SIM would occlude the access of Cys145 to the polyprotein substrate of the protease.

**Telaprevir** (ligand id TEL)

The α-ketoamide group in the best ranking pose of telaprevir is not oriented towards protease residue Cys145, whereas the second-ranking pose (binding energy -8.61) is oriented towards Cys145 (Figure S2). In the latter pose the distance between the S atom of Cys145 and C of TEL is 2.9 Å, which is similar to the distance in the 13b- protease complex. This pose is stabilized hydrogen bonds of TEL with side chain of N^ε^ His41 and backbone N atoms of Gly143, Ser144, Cys145 and Glu166. Thr25 and Gln189 residues of the protease further stabilize the binding of TEL through hydrophobic interactions.

**Vaniprevir** (ligand id VAN)

Docking with vaniprevir provided a lowest scoring pose with predicted binding energy of -10.95 kcal/mol (Figure 1D and Table S1). In this pose (Figure S2), VAN occupies a much a larger portion of the binding cavity than inhibitor 13b. This could be attributed to the large size and macrocyclic nature of VAN. The His41 sidechain N^ε^, Asn142 sidechain O^δ^, Gly143 backbone amide N, and Glu166 backbone amide N atoms form hydrogen bonds with various moieties of VAN. This VAN binding mode includes extensive hydrophobic interactions with the protease, involving residues Thr25, Phe140, Asn142, Glu166 and Pro168.

## References

Anand, K., Palm, G.J., Mesters, J.R., Siddell, S.G., Ziebuhr, J., and Hilgenfeld, R. (2002). Structure of coronavirus main proteinase reveals combination of a chymotrypsin fold with an extra alpha-helical domain. EMBO J 21, 3213–3224. https://doi.org/10.1093/emboj/cdf327

Angelini, M.M., Akhlaghpour, M., Neuman, B.W., and Buchmeier, M.J. (2013). Severe acute respiratory syndrome coronavirus nonstructural proteins 3, 4, and 6 induce double-membrane vesicles. mBio 4. https://doi.org/10.1128/mBio.00524-13

Anson, B., Chapman, M., Lendy, E., Pshenychnyi, S., Richard, T., Satchell, K., and D Mesecar, A. (2020). Broad-spectrum inhibition of coronavirus main and papain-like proteases by HCV drugs Research Square (Preprint). https://doi.org/https://assets.researchsquare.com/files/rs-26344/v1_stamped.pdf

Anson, B., and Mesecar, A. (2020). X-ray structure of SARS-CoV-2 main protease bound to boceprevir at 1.45 A. PDB ID 6WNP:. https://doi.org/10.2210/pdb6WNP/pdb

Bafna, K., Krug, R.M., and Montelione, G.T. (2020). Structural similarity of SARS-CoV2 M(pro) and HCV NS3/4A proteases suggests new approaches for identifying existing drugs useful as COVID-19 therapeutics (Preprint). ChemRxiv. https://doi.org/10.26434/chemrxiv.12153615.v1

Daczkowski, C.M., Dzimianski, J.V., Clasman, J.R., Goodwin, O., Mesecar, A.D., and Pegan, S.D. (2017). Structural insights into the interaction of coronavirus papain-like proteases and Interferon-Stimulated Gene Product 15 from different species. J Mol Biol 429, 1661–1683. https://doi.org/10.1016/j.jmb.2017.04.011

Eastman, R.T., Roth, J.S., Brimacombe, K.R., Simeonov, A., Shen, M., Patnaik, S., and Hall, M.D. (2020). Remdesivir: A review of Its discovery and development leading to emergency use authorization for treatment of COVID-19. ACS Cent Sci 6, 672–683. https://doi.org/10.1021/acscentsci.0c00489

Forli, S., Huey, R., Pique, M.E., Sanner, M.F., Goodsell, D.S., and Olson, A.J. (2016). Computational protein–ligand docking and virtual drug screening with the AutoDock suite. Nature Protocols 11, 905. https://doi.org/10.1038/nprot.2016.051

Fu, L., Ye, F., Feng, Y., Yu, F., Wang, Q., Wu, Y., Zhao, C., Sun, H., Huang, B., Niu, P., et al. (2020). Both Boceprevir and GC376 efficaciously inhibit SARS-CoV-2 by targeting its main protease. Nat Commun 11, 4417. https://doi.org/10.1038/s41467-020-18233-x

Fu, Z., and Huang, H. (2020). SARS CoV-2 PLpro in complex with GRL0617. PDB ID 7CJM. https://doi.org/10.2210/pdb7CJM/pdb

Gorbalenya, A.E., Koonin, E.V., Donchenko, A.P., and Blinov, V.M. (1989). Coronavirus genome: prediction of putative functional domains in the non-structural polyprotein by comparative amino acid sequence analysis. Nucleic Acids Res 17, 4847–4861. https://doi.org/10.1093/nar/17.12.4847

Gordon, D.E., Jang, G.M., Bouhaddou, M., Xu, J., Obernier, K., White, K.M., O’Meara, M.J., Rezelj, V.V., Guo, J.Z., Swaney, D.L., et al. (2020). A SARS-CoV-2 protein interaction map reveals targets for drug repurposing. Nature. https://doi.org/10.1038/s41586-020-2286-9

Grum-Tokars, V., Ratia, K., Begaye, A., Baker, S.C., and Mesecar, A.D. (2008). Evaluating the 3C-like protease activity of SARS-Coronavirus: recommendations for standardized assays for drug discovery. Virus Res 133, 63–73. https://doi.org/10.1016/j.virusres.2007.02.015

Holm, L., and Sander, C. (1993). Protein structure comparison by alignment of distance matrices. J Mol Biol 233, 123–138. https://doi.org/10.1006/jmbi.1993.1489

Holm, L., and Sander, C. (1999). Protein folds and families: sequence and structure alignments. Nucleic Acids Res 27, 244–247. https://doi.org/10.1093/nar/27.1.244

Ianevski, A., Giri, A.K., and Aittokallio, T. (2020). SynergyFinder 2.0: visual analytics of multi-drug combination synergies. Nucleic Acids Res 48, W488–W493. https://doi.org/10.1093/nar/gkaa216

Jin, Z., Du, X., Xu, Y., Deng, Y., Liu, M., Zhao, Y., Zhang, B., Li, X., Zhang, L., Peng, C., et al. (2020). Structure of M(pro) from COVID-19 virus and discovery of its inhibitors. Nature. https://doi.org/10.1038/s41586-020-2223-y

Kester, W.R., and Matthews, B.W. (1977). Comparison of the structures of carboxypeptidase A and thermolysin. J Biol Chem 252, 7704–7710.

Kolb, P., Ferreira, R.S., Irwin, J.J., and Shoichet, B.K. (2009). Docking and chemoinformatic screens for new ligands and targets. Curr Opin Biotechnol 20, 429–436. https://doi.org/10.1016/j.copbio.2009.08.003

Kolb, P., and Irwin, J.J. (2009). Docking screens: right for the right reasons? Curr Top Med Chem 9, 755–770. https://doi.org/10.2174/156802609789207091

Lindner, H.A., Lytvyn, V., Qi, H., Lachance, P., Ziomek, E., and Menard, R. (2007). Selectivity in ISG15 and ubiquitin recognition by the SARS coronavirus papain-like protease. Arch Biochem Biophys 466, 8–14. https://doi.org/10.1016/j.abb.2007.07.006

Ma, C., Sacco, M.D., Hurst, B., Townsend, J.A., Hu, Y., Szeto, T., Zhang, X., Tarbet, B., Marty, M.T., Chen, Y., et al. (2020). Boceprevir, GC-376, and calpain inhibitors II, XII inhibit SARS-CoV-2 viral replication by targeting the viral main protease. Cell Res. https://doi.org/10.1038/s41422-020-0356-z

McGonagle, D., Sharif, K., O’Regan, A., and Bridgewood, C. (2020). The role of cytokines including Interleukin-6 in COVID-19 induced pneumonia and macrophage activation syndrome-like disease. Autoimmun Rev 19, 102537. https://doi.org/10.1016/j.autrev.2020.102537

Morris, G.M., Huey, R., Lindstrom, W., Sanner, M.F., Belew, R.K., Goodsell, D.S., and Olson, A.J. (2009). AutoDock4 and AutoDockTools4: Automated docking with selective receptor flexibility. J Comput Chem 30, 2785–2791. https://doi.org/10.1002/jcc.21256

Orengo, C.A., Michie, A.D., Jones, S., Jones, D.T., Swindells, M.B., and Thornton, J.M. (1997). CATH--a hierarchic classification of protein domain structures. Structure 5, 1093–1108. https://doi.org/10.1016/s0969-2126(97)00260-8

Oudshoorn, D., Rijs, K., Limpens, R., Groen, K., Koster, A.J., Snijder, E.J., Kikkert, M., and Barcena, M. (2017). Expression and cleavage of Middle East Respiratory Syndrome Coronavirus nsp3-4 polyprotein induces the formation of double- membrane vesicles that mimic those associated with coronaviral RNA replication. mBio 8. https://doi.org/10.1128/mBio.01658-17

Ozen, A., Prachanronarong, K., Matthew, A.N., Soumana, D.I., and Schiffer, C.A. (2019). Resistance outside the substrate envelope: hepatitis C NS3/4A protease inhibitors. Crit Rev Biochem Mol Biol 54, 11–26. https://doi.org/10.1080/10409238.2019.1568962

Pan, H., Peto, R., Abdool Karim, Q., Alejandria, M., HenaoRestrepo, A.M., Hernández García, C., Kieny, M.-P., Malekzadeh, R., Murthy, S., Preziosi, M.-P., et al. (2020). Repurposed antiviral drugs for COVID-19 – interim WHO SOLIDARITY trial results (preprint). MedRciv. https://doi.org/10.1101/2020.10.15.20209817

Peng, Q., Peng, R., Yuan, B., Zhao, J., Wang, M., Wang, X., Wang, Q., Sun, Y., Fan, Z., Qi, J., et al. (2020). Structural and biochemical characterization of the nsp12- nsp7-nsp8 core polymerase complex from SARS-CoV-2. Cell Rep 31, 107774. https://doi.org/10.1016/j.celrep.2020.107774

Robertus, J.D., Alden, R.A., Birktoft, J.J., Kraut, J., Powers, J.C., and Wilcox, P.E. (1972). An x-ray crystallographic study of the binding of peptide chloromethyl ketone inhibitors to subtilisin BPN’. Biochemistry 11, 2439–2449. https://doi.org/10.1021/bi00763a009

Sheahan, T.P., Sims, A.C., Zhou, S., Graham, R.L., Pruijssers, A.J., Agostini, M.L., Leist, S.R., Schafer, A., Dinnon, K.H., 3rd, Stevens, L.J., et al. (2020). An orally bioavailable broad-spectrum antiviral inhibits SARS-CoV-2 in human airway epithelial cell cultures and multiple coronaviruses in mice. Sci Transl Med 12. https://doi.org/10.1126/scitranslmed.abb5883

Snijder, E.J., Limpens, R., de Wilde, A.H., de Jong, A.W.M., Zevenhoven-Dobbe, J.C., Maier, H.J., Faas, F., Koster, A.J., and Barcena, M. (2020). A unifying structural and functional model of the coronavirus replication organelle: Tracking down RNA synthesis. PLoS Biol 18, e3000715. https://doi.org/10.1371/journal.pbio.3000715

Swaim, C.D., Canadeo, L.A., Monte, K.J., Khanna, S., Lenschow, D.J., and Huibregtse, J.M. (2020). Modulation of extracellular ISG15 signaling by pathogens and viral effector proteins. Cell Rep 31, 107772. https://doi.org/10.1016/j.celrep.2020.107772

Swaim, C.D., Scott, A.F., Canadeo, L.A., and Huibregtse, J.M. (2017). Extracellular ISG15 signals cytokine secretion through the LFA-1 integrin receptor. Mol Cell 68, 581–590 e585. https://doi.org/10.1016/j.molcel.2017.10.003

Wolff, G., Limpens, R., Zevenhoven-Dobbe, J.C., Laugks, U., Zheng, S., de Jong, A.W.M., Koning, R.I., Agard, D.A., Grunewald, K., Koster, A.J., et al. (2020a). A molecular pore spans the double membrane of the coronavirus replication organelle. Science 369, 1395–1398. https://doi.org/10.1126/science.abd3629

Wolff, G., Melia, C.E., Snijder, E.J., and Barcena, M. (2020b). Double-membrane vesicles as platforms for viral replication. Trends Microbiol 28, 1022–1033. https://doi.org/10.1016/j.tim.2020.05.009

Wu, F., Zhao, S., Yu, B., Chen, Y.M., Wang, W., Song, Z.G., Hu, Y., Tao, Z.W., Tian, J.H., Pei, Y.Y., et al. (2020). A new coronavirus associated with human respiratory disease in China. Nature 579, 265–269. https://doi.org/10.1038/s41586-020-2008-3

Zhang, L., Lin, D., Kusov, Y., Nian, Y., Ma, Q., Wang, J., von Brunn, A., Leyssen, P., Lanko, K., Neyts, J., et al. (2020a). alpha-Ketoamides as broad-spectrum inhibitors of Coronavirus and Enterovirus replication: Structure-based design, synthesis, and activity assessment. J Med Chem. https://doi.org/10.1021/acs.jmedchem.9b01828

Zhang, L., Lin, D., Sun, X., Curth, U., Drosten, C., Sauerhering, L., Becker, S., Rox, K., and Hilgenfeld, R. (2020b). Crystal structure of SARS-CoV-2 main protease provides a basis for design of improved alpha-ketoamide inhibitors. Science 368, 409–412. https://doi.org/10.1126/science.abb3405

Zhao, C., Sridharan, H., Chen, R., Baker, D.P., Wang, S., and Krug, R.M. (2016). Influenza B virus non-structural protein 1 counteracts ISG15 antiviral activity by sequestering ISGylated viral proteins. Nat Commun 7, 12754. https://doi.org/10.1038/ncomms12754

## STAR Methods References

Amanat, F., White, K.M., Miorin, L., Strohmeier, S., McMahon, M., Meade, P., Liu, W.C., Albrecht, R.A., Simon, V., Martinez-Sobrido, L., et al. (2020). An In Vitro Microneutralization Assay for SARS-CoV-2 Serology and Drug Screening. Curr Protoc Microbiol 58, e108. https://doi.org/10.1002/cpmc.108

DeLano, W.L. (2009). The PyMOL Molecular Graphics System, Version 1.2r3pre, Schrödinger, LLC.

Lanevski, A., Giri, A.K., and Aittokallio, T. (2020). SynergyFinder 2.0: visual analytics of multi-drug combination synergies. Nucleic Acids Res 48, W488–W493. https://doi.org/10.1093/nar/gkaa216

Morris, G.M., Goodsell, D.S., Halliday, R.S., Huey, R., Hart, W.E., Belew, R.K., and Olson, A.J. (1998). Automated docking using a Lamarckian genetic algorithm and an empirical binding free energy function. J Comput Chem 19, 1639–1662. https://doi.org/10.1002/(SICI)1096-987X(19981115)19:14<1639::AID-JCC10>3.0.CO;2-B

Salentin, S., Schreiber, S., Haupt, V.J., Adasme, M.F., and Schroeder, M. (2015). PLIP: fully automated protein-ligand interaction profiler. Nucleic Acids Res 43, W443–447. https://doi.org/10.1093/nar/gkv315

Sanner, M.F. (1999). Python: a programming language for software integration and development. J Mol Graph Model 17, 57–61.

Zhang, L., Lin, D., Sun, X., Curth, U., Drosten, C., Sauerhering, L., Becker, S., Rox, K., and Hilgenfeld, R. (2020). Crystal structure of SARS-CoV-2 main protease provides a basis for design of improved alpha-ketoamide inhibitors. Science 368, 409–412. https://doi.org/10.1126/science.abb3405

